# RAD51 promotes non-homologous genetic rearrangements that are prevented by 53BP1

**DOI:** 10.1101/768788

**Authors:** Josée Guirouilh-Barbat, Wei Yu, Loelia Babin, Elisa Yaniz Galende, Tatiana Popova, Ayeong So, Chloé Lescale, Marc-Henri Stern, Erika Brunet, Ludovic Deriano, Bernard S. Lopez

## Abstract

Homologous recombination (HR), which requires long sequence homologies, is considered a high fidelity mechanism, preserving genome stability. In contrast, we show here that the central HR players RAD51 or BRCA2, promote genetic instability, fostering translocations and capture of ectopic chromosomal sequences when joining distant DNA breaks. Surprisingly, these events do not involve sequence homologies. Moreover, our data reveal that 53BP1 protects against RAD51-mediated non-homologous genetic rearrangements. Finally, analysis of a large panel of breast tumors revealed that BRCA2 proficiency is associated with increased frequency of capture of non-homologous sequences at junctions of structural variants (translocations, duplications, inversions, deletions). These data reveal that HR proteins (RAD51, BRCA2) possess the intrinsic capacity to generate genetic instability through sequence homology-independent processes, and that 53BP1 protects against it. We propose that BRCA2/RAD51-mediated genome instability occurs in the course of sequence homology search for HR.

## Introduction

Genetic instability is associated with tumorigenesis, developmental syndromes and pre-mature ageing (Garinis et al., 2008; Hoeijmakers, 2009; Lans and Hoeijmakers, 2006; Negrini et al., 2010). Cells developed elaborate systems protecting genome stability, albeit allowing genetic diversity. While mechanisms involved in genome stability/diversity maintenance are largely documented, the molecular mechanisms driving genetic instability are far from being fully characterized, at a molecular level.

DNA double-strand break (DSB) is one of the most harmful lesions and a major source of genetic variability. Indeed, DSBs generate both genome rearrangements (e.g., translocations, inversions, duplications, deletions, and complex rearrangements) and mutagenesis at the sealed junctions themselves. Cells use two primary strategies to repair DSBs. The first strategy joins and ligates two DNA double-strand ends without requiring sequence homology and is referred to as non-homologous end-joining (NHEJ). The second strategy relies on the use of a DNA partner sharing long sequence homologies and is thus referred to as homologous recombination (HR).

HR, which requires hundreds of base pairs of homology between the two interacting DNA partners, is generally considered a high fidelity process essential for genome stability maintenance. HR is an evolutionarily highly conserved process that can be divided into sub-classes all initiated by common mechanisms (Haber, 2014): 1) Resection of DSBs, generating 3’ single-stranded DNA tails (ssDNA); 2) loading of RAD51 on the ssDNA by BRCA2/PALB2, generating the actual active species of HR, i.e., the ssDNA/RAD51 filament; 3) search for sequence homologies and strand invasion of a homologous intact duplex DNA, representing thus the pivotal steps of HR performed by the ssDNA/RAD51 filament; and 4) DNA synthesis primed on the invading 3’ end. Then, depending on the subsequent steps and resolution of intermediates, the process will lead to gene conversion associated or not with crossovers, to break–induced replication (BIR) or to synthesis-dependent strand annealing (SDSA). Importantly, the length of homology between the two partners is a paramount parameter for HR. This point was extensively studied at the end of the last millennium leading to the definition of the MEPS (minimal effective treatment segment), ranging between 250 and 300 bp in mammalian cells (Bollag et al., 1989; Liskay et al., 1987; Lopez et al., 1992; Rubnitz and Subramani, 1984). Therefore, it is widely admitted that HR requires hundred(s) bp of sequence homology, in mammalian cells.

In addition to HR and NHEJ (in fact the KU-ligase IV-dependent canonical NHEJ, C-NHEJ), alternative processes mediated by microhomologies (2-4 bp) have been described: alternative end-joining (A-EJ), also called backup NHEJ (B-NHEJ) or microhomology-mediated end-joining (MMEJ); microhomology-mediated BIR (MMBIR); fork stalling and template switching (FoSTeS); and microhomology-mediated template switching (MMTS) (for review see (Deriano and Roth, 2013; So et al., 2017)). Because they are independent of KU-ligase 4 and they do not require long sequence homologies, but only few bp of microhomology (thus far from the MEPS), these events are distinct from both C-NHEJ and HR.

Using an intrachromosomal substrate (CD4-3200bp), we have shown that few kb of distance between two DNA double-stranded ends are sufficient to favor the capture of non-homologous ectopic chromosomal sequences (≥ 45 bp) at the sealed junction. Such kind of insertions jeopardizes genome integrity and is commonly observed in cancer cells. However, the underlying mechanisms remain to be characterized. The borders of the insertions are characterized by the absence of homologous sequences and, in contrast, the presence of microhomologies or unmappable patches of nucleotides; therefore, these events do not correspond to HR-related events (Guirouilh-Barbat et al., 2016), but rather to microhomology-mediated insertions. We proposed that MMTS mediates the captures of ectopic sequences (Guirouilh-Barbat et al., 2016). The cohesion complex, which restrains the joining of distant ends by channeling DSB repair toward the use of sister chromatids, also limits the capture of ectopic sequences (Gelot et al., 2016). Because sister chromatid exchanges involve HR, we postulated that ablation of HR should also stimulate ectopic sequences capture. Surprisingly, silencing the HR proteins RAD51 or BRCA2 did not stimulate ectopic sequences capture but instead decreased it, although no homologous sequences were involved. The impact of RAD51 depletion was more particularly observed in cells depleted of 53BP1 in which ectopic sequences capture is favored. In addition, we found that RAD51 was also implicated in the generation of inter-chromosomal translocations themselves fostered by 53BP1 depletion, although no sequence homology was involved. Therefore the pivotal HR player RAD51 generates genetic instability in a process that does not require long sequence homology. This activity strongly contrasts with the canonical role of RAD51 in HR. In addition, our data reveal that 53BP1 protects against RAD51-mediated genetic instability. We propose a model in which BRCA2/RAD51 promote genome instability through non-homologous recombination (or micro-homologous recombination) when the RAD51/ssDNA filament scans the DNA in the course of homology search for HR. These data unravel a molecular mechanism driving genetic instability: mediated by the pivotal HR players RAD51 and counteracted by 53BP1. Therefore a balance between RAD51 and 53BP1 activities governs genome stability maintenance.

## Results

### 53BP1 counteracts the capture of non-homologous ectopic sequences promoted by RAD51 and BRCA2, at junctions of distant DSBs

Using the intrachromosomal substrate CD4-3200bp (Fig. 1A), we previously reported the capture of ectopic sequences at the joining of 3200 bp distant ends, in a sequence homology-independent manner. Moreover, we show that ablation of 53BP1 strongly increased the frequency of ectopic non-homologous sequences capture (Guirouilh-Barbat et al., 2016). Here we show that silencing the pivotal HR protein RAD51 or BRCA2 completely abolished the stimulation of sequence capture resulting from 53BP1 silencing (Fig. 1B-D, Table 1 and Supplementary data S1). These results were unexpected because the CD4-3200bp substrate does not share sequence homology with the human genome, and thus ectopic sequence capture cannot be scored as HR events. In the presence of 53BP1, ablation of RAD51 or BRCA2 barely affected the frequency of ectopic sequences capture (Supplementary data S2).

**Fig. 1:**
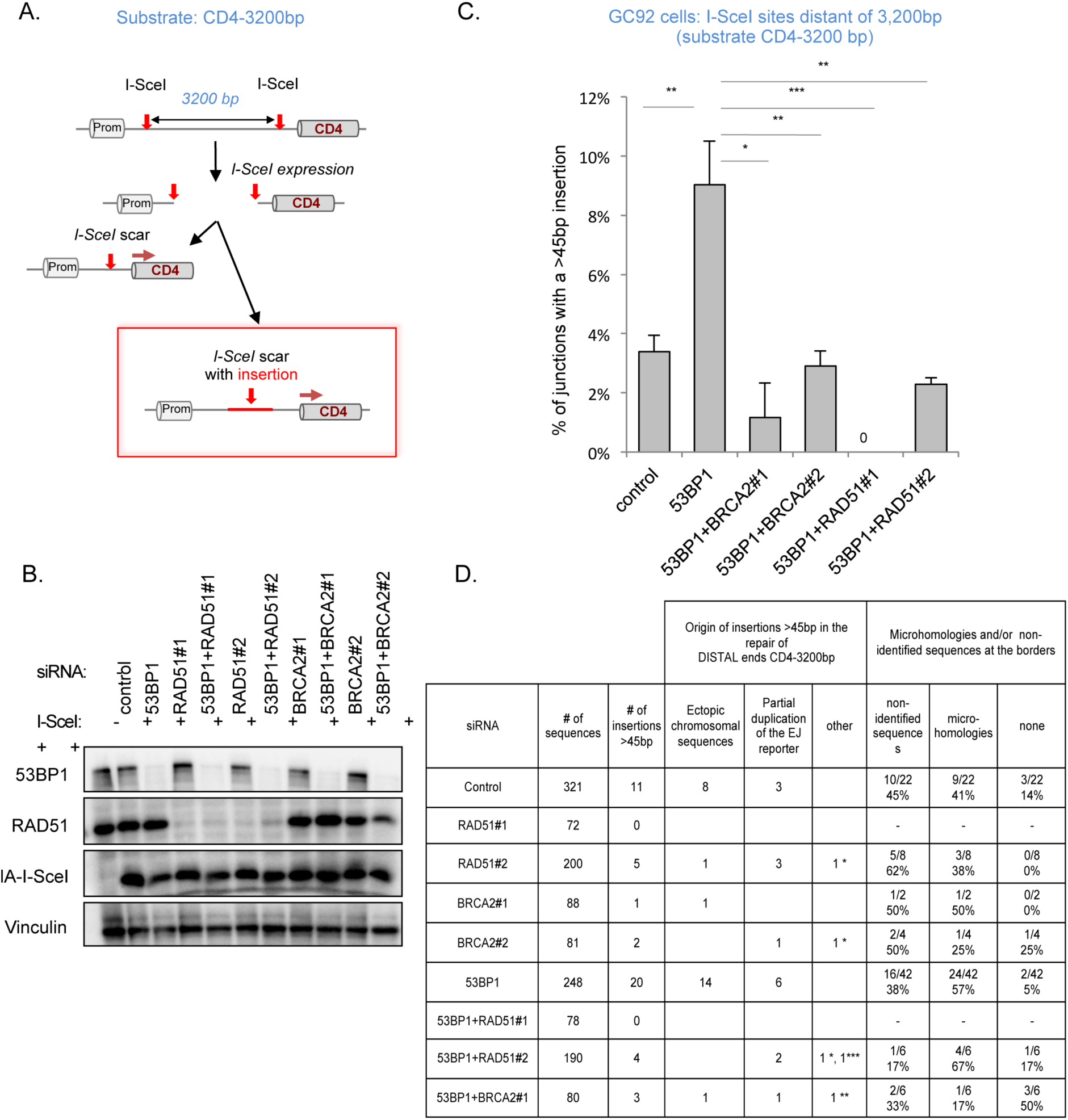
BRCA2/RAD51 are required for ectopic sequences capture at the junction scars of distant ends. **A.** The CD4-3200bp substrate. **B.** Analysis of protein depletion and I-SceI expression in GC92 cells. **C.** Percentage of junctions (from I-SceI sites, 3200bp apart) containing ectopic sequences capture in the GC92 cells after 53BP1, RAD51 and/or BRCA2 depletion. Histograms represent the percentage of junctions with insertions >45 bp. Each value is the average of 2 to 7 independent experiments ± SEM. **D.** Origin of large insertions (>45bp) monitored in the repair of distal DSEs (CD4-3200bp) upon 53BP1, RAD51 and/or BRCA2 depletion, in the GC92 cell line. Values are calculated from 2 to 7 independent experiments and sequencing of 72 to 321 junction sequences (* I-SceI expression plasmid, ** bovine satellite, *** the whole insertion is non mappable).

Some captured sequences were partial duplications of the intrachromosomal reporter substrate, but the majority originated from other chromosomes (Fig. 1D and Supplementary data S1). Although both intra- and inter-chromosomal sequences captures are RAD51/BRCA2 dependent, no long stretches of sequence homology were present at the borders (Supplementary data S1); instead, 2-4 bp microhomologies or unmappable stretches of nucleotides were present. Few borders exhibited neither microhomologies nor unmappable sequences (Fig. 1D).

Therefore, in contrast with the canonical sequence homology-dependent role of RAD51/BRCA2 in HR and genome stability maintenance, RAD51 and BRCA2 promote the capture of non-homologous ectopic sequences, in human cells. In addition, these data reveal that 53BP1 protects against this RAD51-promoted genetic instability.

### Genome-wide analysis of chromosomal rearrangements at the I-­SceI site

We then assessed the impact of RAD51 on genetic rearrangements at one DSB, at the genome wide level. We used linear amplification-mediated high-throughput genome-wide translocation sequencing (LAM-HTGTS) (Frock et al., 2015; Hu et al., 2016) to detect genome-wide chromosomal rearrangements at the CD4-3200bp chromosomal locus. As a bait for LAM-HTGTS, we chose the I-SceI DSB site close to the CD4 promoter of the CD4-3200bp chromosomal substrate and used two primers located 320 bp and 139 bp 5’ upstream of the I-SceI site to amplify breakpoint junction sequences (Fig. 2A).

**Fig. 2:**
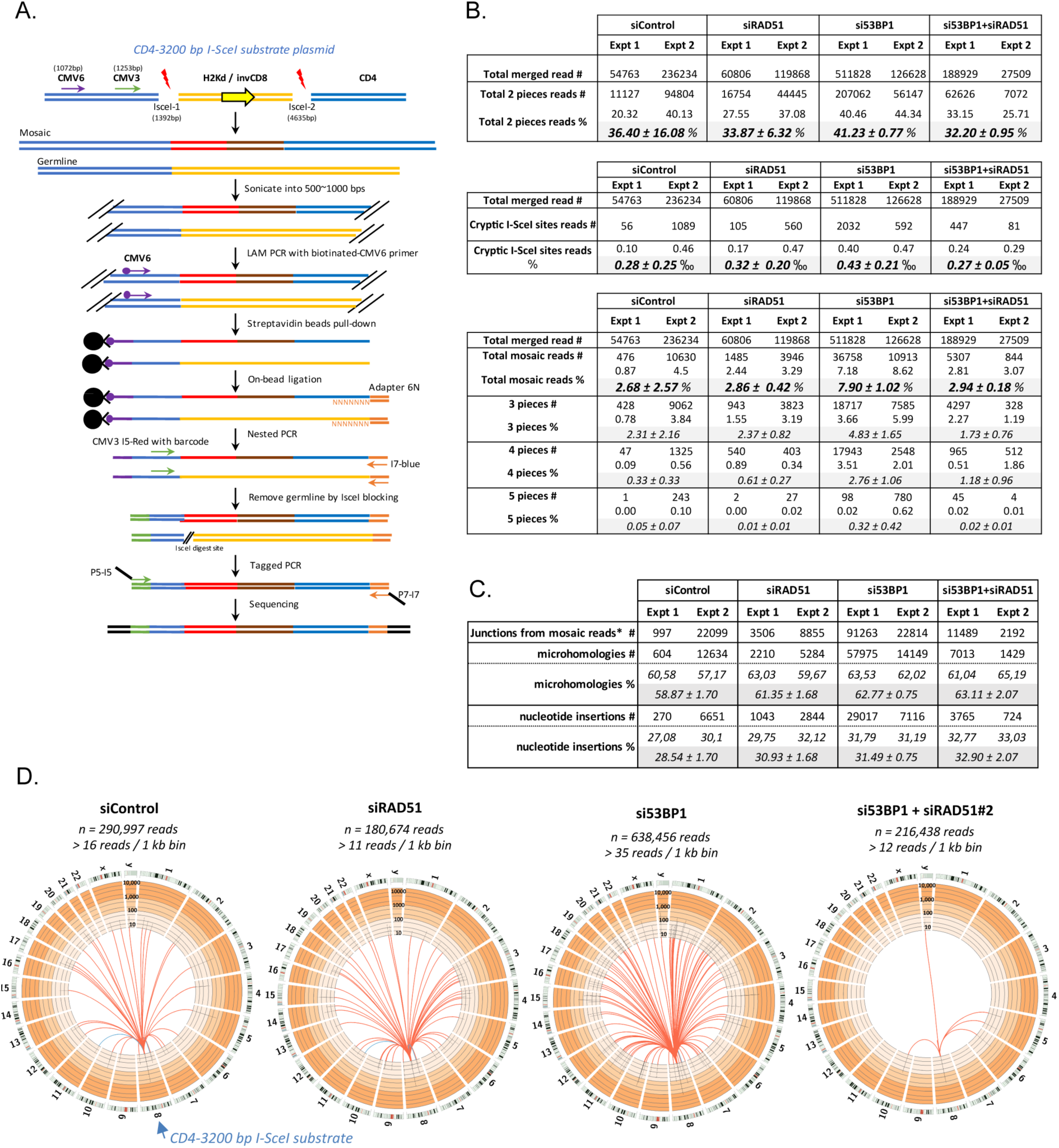
Linear amplification-mediated high-throughput genome-wide translocation sequencing (LAM-HTGTS) **A.** Linear amplification-mediated high-throughput genome-wide translocation sequencing (LAM-HTGTS) methodology outline. **B.** Numbers (#) of sequences containing 2 identified (upper table), I-SceI cryptic sites (middle table) or more than two identified pieces (lower table) and their percentages (%) relative to the total number of merged reads in control or 53BP1, RAD51 and 53BP1/RAD51-depleted GC92 cells. The results from two independent experiments are shown. **C.** Microhomology usage and presence of non-identified patches of nucleotides at junctions between identified pieces in mosaic reads containing 3 and 4 pieces. Numbers (#) and percentages (%) for each sample are indicated. The results from two independent experiments are shown. **D.** Circos plots representing interchromosomal junctions in control, RAD51- or 53BP1-depleted and 53BP1/RAD51-depleted GC92 cells. The CD4-3200bp substrate is located on chromosome 8 (blue arrow). The number of total merged reads for two combined experiments and the cutoff of reads for 1 kb bins are indicated. The cutoff of reads is proportional to the total merged reads. The bins with a number of captured reads superior to the cutoff are linked by red or blue lines (red line: CD4-3200bp substrate-human genome joins; blue line: CD4-3200bp substrate-human genome with cryptic I-SceI site joins). The numbers of joins are plotted as black bars on a log scale with the indicated custom axis from 10 to 10000.

We analyzed genome-wide rearrangements at this I-SceI site upon I-SceI expression and silencing either RAD51, 53BP1 or both 53BP1 and RAD51. Analysis of the sequencing reads confirmed that a single copy of the CD4-3200bp substrate was located on chromosome 8 at approximately position 76,367 kb. We next searched for the presence of split reads among merged reads (Supplementary Fig. S3A). As expected, a majority of discordant reads corresponded to two-pieces split reads between the 5’ promoter sequence of the I-SceI cutting site and sequences within the CD4-3200 bp substrate (Supplementary Fig. S3B). These events mainly correspond to end-joining events, as we have shown before (Grabarz et al., 2013; Guirouilh-Barbat et al., 2004, 2007; Rass et al., 2009). Other two-pieces split reads contained the bait promoter sequence and either unmappable sequences, the I-SceI donor plasmid sequence or sequences mapped to the human genome, including chromosomal translocations (Supplementary Fig. S3B). This latter category might also result from the direct end-joining of I-SceI-induced DSBs to prey DSBs occurring elsewhere in the genome.

Consistent with this, we identified multiple sequences located at or near previously described cryptic I-SceI target sites (Frock et al., 2015), which likely represent direct end-joining events between two I-SceI-induced DSBs (Fig. 2B and Supplementary Fig. S3B). Therefore, two-pieces split reads correspond to rearrangements/translocations occurring by a mix of end-joining, and/or microhomology mediated mechanisms. These events were more frequent in 53BP1-depleted cells but this increase was abolished by co-depletion of RAD51 (Fig. 2B), showing that 1-RAD51 is implicated in some of microhomologies mediated rearrangements and 2-53BP1 protects against these RAD51-mediated events.

In spite of the relatively short length of the reads (250-450bp) we also detected mosaic reads containing multiple (>2) mappable parts (Fig. 2B and Supplementary Fig. S3B). Most mosaics contained 3 mappable pieces, although four and even five pieces were occasionally recorded at lower frequencies (Fig. 2B and Supplementary Fig. S3B). Consistent with our previous analysis (see Fig. 1), silencing 53BP1 increased the occurrence of mosaics by approximately 3-fold compared to wild-type cells (Fig. 2B). Silencing RAD51 in 53BP1-silenced cells abolished the stimulation of such events (Fig. 2B).

Approximately 60% of the breakpoint junctions in mosaic reads containing 3 and 4 pieces exhibited microhomologies (Fig. 2C), with a median of 3 nucleotides in control cells and up to 6 bp in 53BP1-silenced cells (Supplementary Fig. S4), and nearly 30% of joints contained non-identified nucleotide insertions (Fig. 2C), with a median of 9bp in control cells up to 20bp in 53BP1-silenced cells (Supplementary Fig. S4). These percentages are similar to those obtained in our previous and present experiments with the CD4-3200 bp substrate (Fig. 1, Supplementary data S1 and (Guirouilh-Barbat et al., 2016). Only few junctions contained neither microhomologies nor non-identified nucleotide insertions (Fig. 2C).

A large fraction of mosaic reads contained DNA sequences that mapped to the human genome (Fig. 2D and Supplementary Fig. S5), thus representing the capture of human ectopic chromosomal sequences or translocations. As the cryptic I-SceI sites were not over-represented in mosaic events, such events arise more likely through MMTS rather than direct DSB End-Joining. These mosaic events were stimulated in 53BP1-silenced cells but abolished in 53BP1- and RAD51-silenced cells (Fig. 2D and Supplementary Fig. S5).

These analyses show that RAD51 generates both genomic rearrangements themselves and capture of ectopic sequences at the junctions, in a sequence-homology- independent manner, and that 53BP1 counteracts RAD51-mediated non-homologous rearrangements.

### 53BP1 protects against RAD51-mediated t(2;5) translocation between non-homologous sequences

As LAM-HTGTS suggested that RAD51 fosters translocations independently of sequence homology, and that this is prevented by 53BP1, we directly addressed the impact of 53BP1 and RAD51 on the occurrence of a sequence homology-independent translocation targeted by CRISPR-Cas9, at another locus and in another cell line. The t(2;5) translocation between ALK and NPM1 sequences occurs in anaplasic large cell lymphoma (ALCL). Such translocation can be reconstructed in cultured cells using specific CRISPR-Cas9 targeting (Piganeau et al., 2013) (Fig. 3A). Neither 53BP1 nor RAD51 silencing (Fig. 3B) significantly affected the efficiency of Cas9-GFP expression (Supplementary Fig. S6A) and the efficiency of cleavage by Cas9 monitored by TIDE analysis of INDELS at both sites (Supplementary Fig. S6B).

**Fig. 3.**
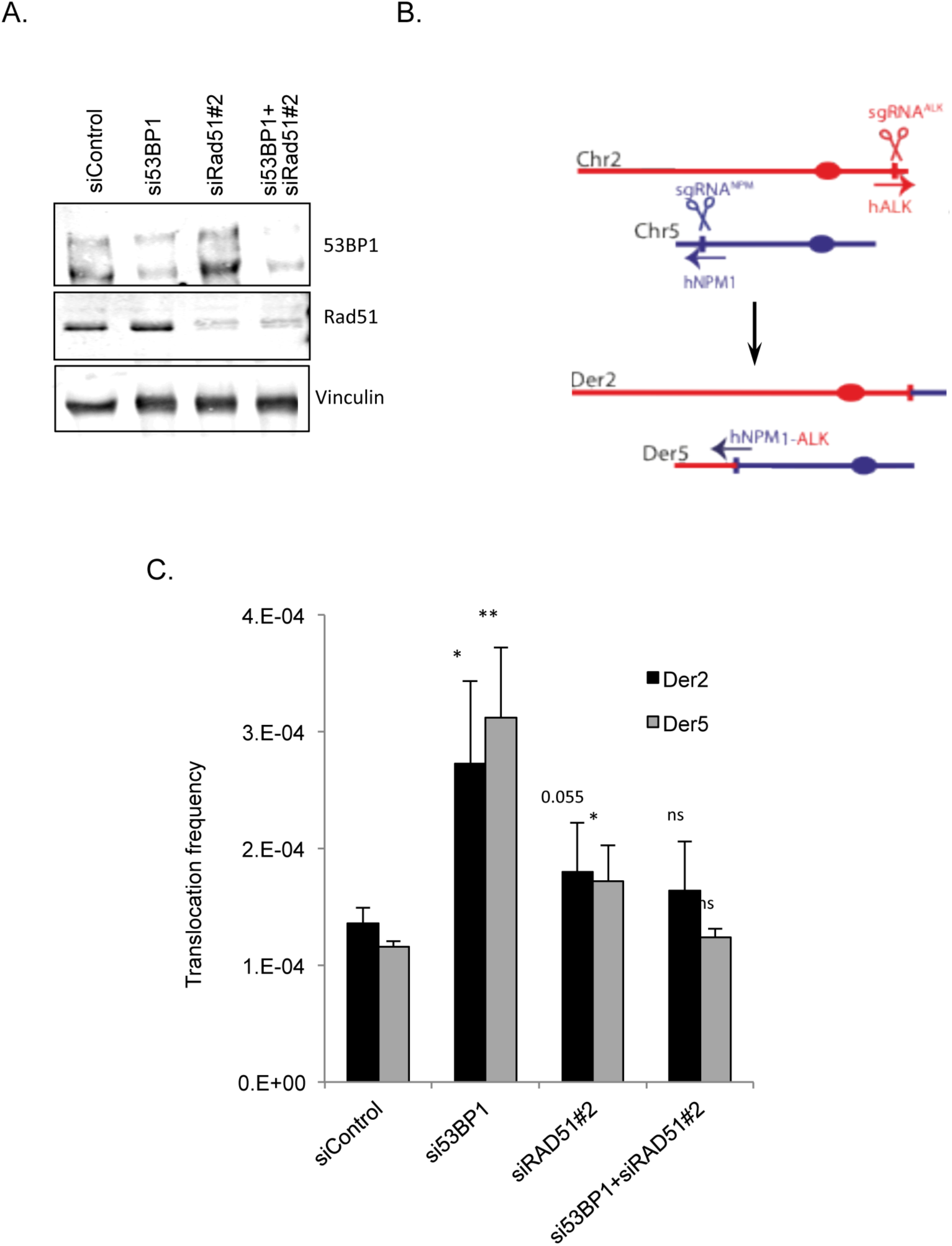
The t(2;5) translocation between ALK and NPM1 induced by specific CRISPR-Cas9. **A.** Scheme of the reconstituted t(2;5) translocation with the 2 derivative chromosomes (der2 and der5). **B.** Analysis of 53BP1 and Rad51 expression. **C.** Frequency of CRISPR-CAS9-induced t(2;5) hALK/hNPM1 translocation. The data correspond to the mean and SEM of 3 independent experiments.

In this novel set up, ablation of 53BP1 also resulted in a strong increase in the t(2;5) translocation frequency (Fig. 3C). Moreover, although no sequence homology was involved, ablation of RAD51 abolished the stimulation of translocation resulting from 53BP1 silencing (Fig. 3C). These data are fully consistent with that obtained with the LAM-HTGTS analysis described above, at a different locus, in a different cell line.

Collectively, our data point RAD51 as a driver of genetic instability, through events that do not involve sequence homology and 53BP1 as a protector against such events.

### Capture of ectopic sequences occurs at SV junctions in human breast tumors

Then, we analyzed the junction scars of structural variants (SVs), *i.e*. translocations, duplications, deletions and inversions, in a physiopathological context. We considered the set of SV junctions from the whole genome sequencing (WGS) of 560 breast cancers described previously by S.Nik-Zainal et al (Nik-Zainal et al., 2016). For the basal analysis of junctions in breast tumors, we first excluded the cases with HR deficiency (HRD tumors) in which BRCA1/2 are inactivated or predicted to be inactivated (see M&Ms), in order to avoid any bias associated with HR deficient phenotype. This gave 419 cases corresponding to 47833 SVs, i.e., translocations, duplications, inversions, or deletions, therefore involving distant partners (>1000bp). 63% of SV junctions exhibited micro-homologies (Supplementary data Fig. S7), which is very similar to the frequency observed in the above HTGTS analysis (Fig. 2C). Scar junctions contained sequence insertions in 19% of SVs, including 3% of junctions with insertions of at least 45 bp that we have defined as the cut-off size of ectopic sequences capture (Guirouilh-Barbat et al., 2016) (Fig. 4A); this is in similar range than that observed with the CD4-3200 bp substrate (compare Fig. 4A and 4B). Of note, each different type of SV (namely, duplications, inversions, deletions, and translocations) exhibited similar frequencies of ectopic sequences capture at the scar junctions (Fig. 4A and Supplementary data Fig. S6).

**Fig. 4.**
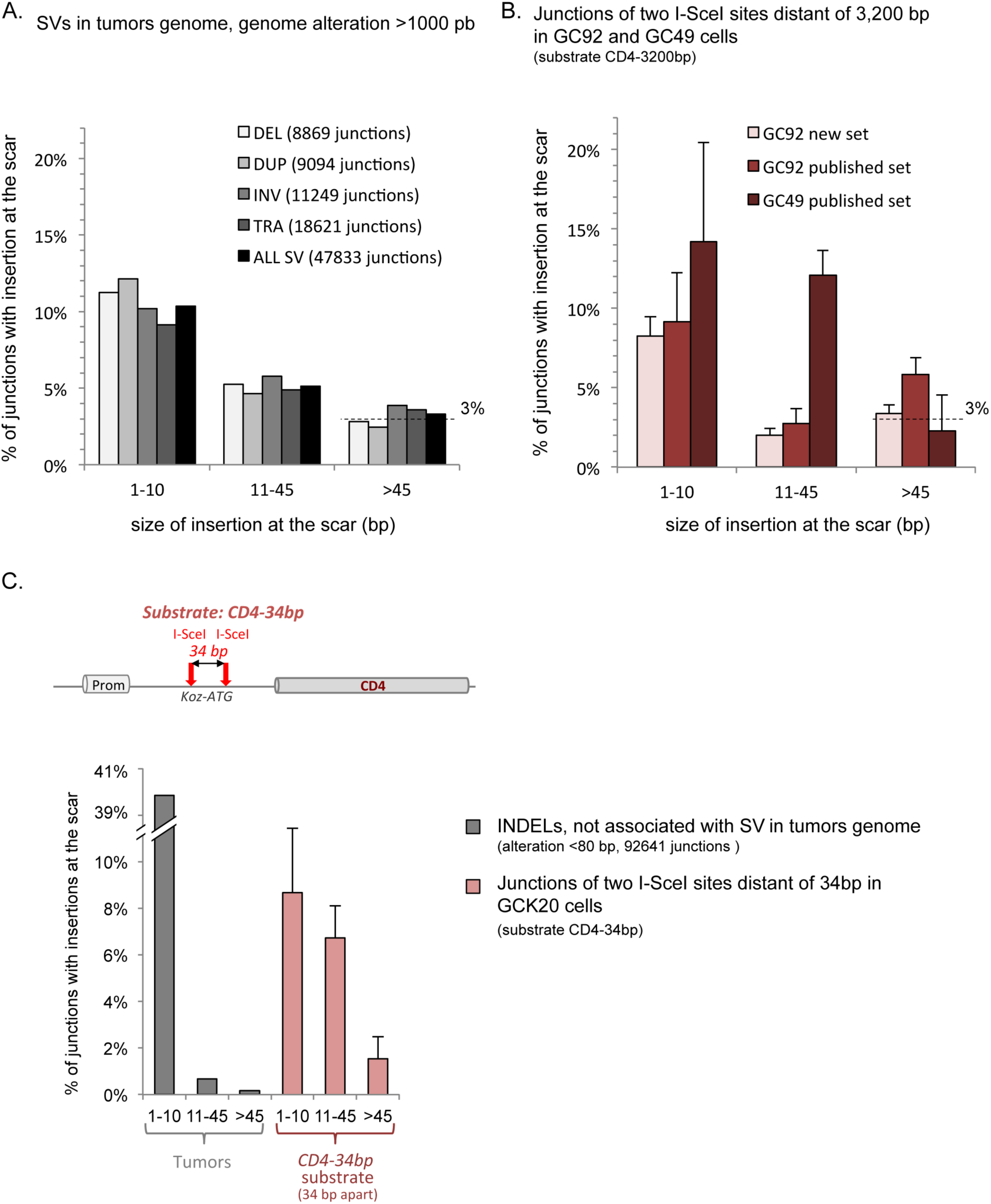
Capture of ectopic sequences at the junction of SVs in breast tumors. **A.** Frequency of insertions at the junction scars (total # of analyzed junctions: 47833) of SVs (size of alteration > 1,000 bp) in a panel of 419 non HRD breast tumors. Approximately 20% of junctions contain insertions in all types of SVs. Histograms represent the fraction of scars with insertions of 1-10 bp, 11-45 bp or >45 bp; we propose that the latter arise through MMTS (Guirouilh-Barbat et al., 2016). **B.** Frequency of insertions at the junction scars of two I-SceI breakage sites 3,200 bp apart (substrate CD4-3200 bp, see Fig. 1B). GC92 and GC49 are two different clones bearing the CD4-3200 bp reporter (Guirouilh-Barbat et al., 2016). The new set of data represent the average of 5-7 independent experiments ± SEM. Published data were from (Guirouilh-Barbat et al., 2016). **C.** Gray bars: Frequency of insertions at the junction scars of INDELs not associated with SVs in tumors (size of alteration < 80 bp, total # of analyzed INDELs: 92641). Red bars: Frequency of insertions at the junction scars of two I-SceI breakage sites 34 bp apart in GCK20 cells. The substrate (CD4-34 bp) used is shown in the upper panel (Guirouilh-Barbat et al., 2016). The results represent the average of 5 independent experiments ± SEM that were used in a previous publication (Guirouilh-Barbat et al., 2016).

No long homologous sequences were present at the borders of the ectopic sequences capture events, suggesting that they occur through non-homologous-mediated process(es). Instead, the borders exhibit microhomologies and non-identified sequences, similar to those observed with the CD4-3200 bp intrachromosomal substrate.

We also analyzed the INDELs (small genome alteration < 80bp) not associated with SVs (92641 junctions), therefore that do not involve rearrangements of distant DNA partners. Insertions >45bp are only barely detectable (0.17% of all insertions) (Fig. 4C). This corresponds to the experimental assay with CD4-34 bp intra-chromosomal substrates (a derivative of the CD4-3200 bp where the two DNA ends are separated by only 34 bp) showing that the joining of close double-strand ends does not permit the capture of ectopic sequences (Fig. 4C and (Guirouilh-Barbat et al., 2016)). Taken together, the data confirm that capture of ectopic sequences is favored at junctions of distant partners, in non-HRD breast tumors.

### Captures of non-homologous ectopic sequences at SV junctions are decreased in BRCA2-deficient tumors

The cohort of breast tumors also contains 28 cases with BRCA2 inactivation due to germline or somatic deleterious mutations and loss of the second allele (Nik-Zainal et al., 2016). Thus, we compared BRCA2 deficient tumors to the control group (high grade HER2-negative cases without other HR deficiency) for the frequency of non-homologous sequence capture at the SVs junctions (Fig. 5).

**Fig. 5.**
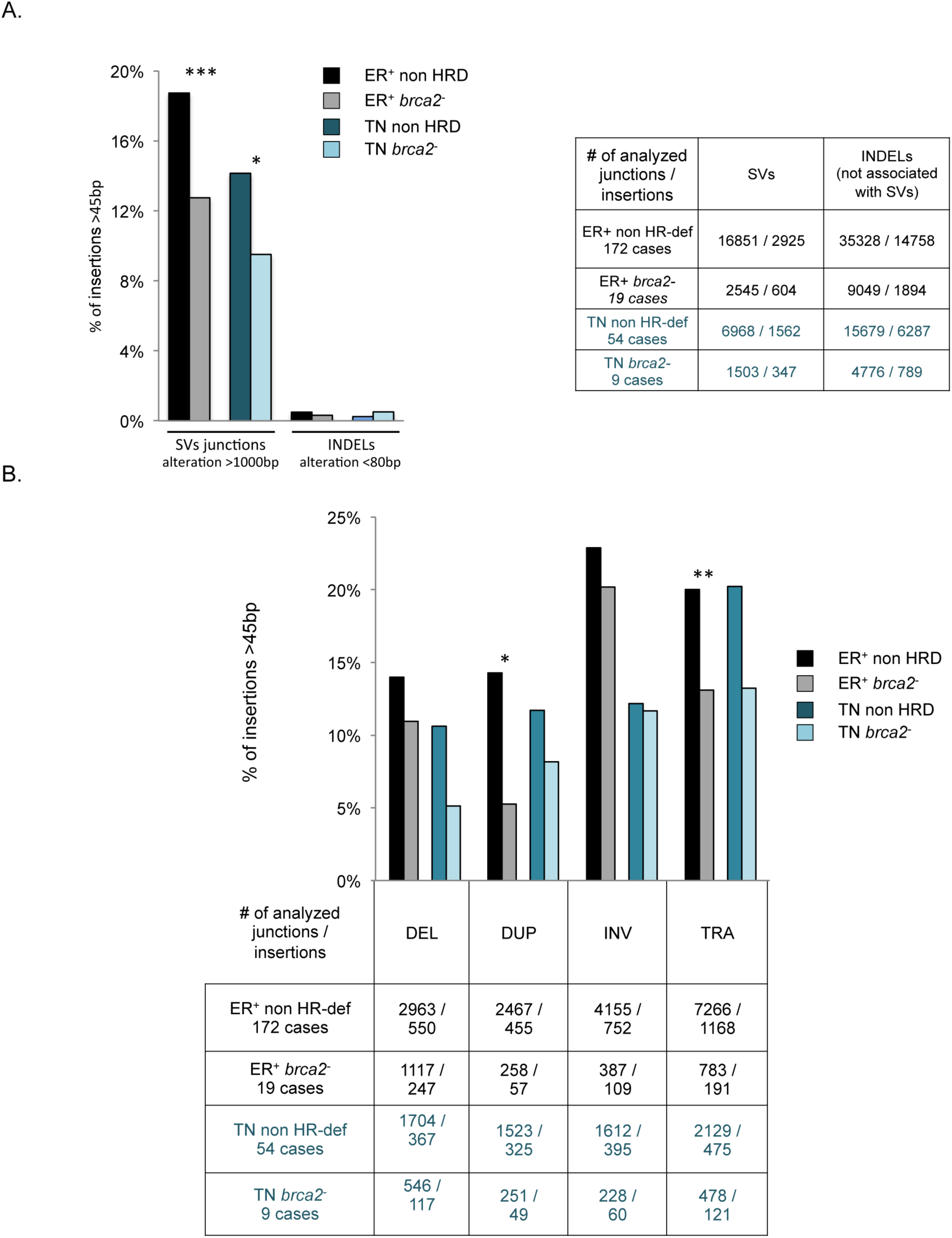
Capture of ectopic sequences at SV scars in human non HR-deficient (non HR-def) *versus BRCA2*-deficient breast tumors. **A.** Ectopic sequences capture in total SVs versus INDELs in non-SV events. Left four bars: SVs, involving distant DNA. Right four bars: ectopic sequences capture at INDELs not associated with SVs, i.e. that do not involve distant DNA. **B.** Ectopic sequences capture in each kind of SVs. The bars represent the percentage of insertions that are >45 bp among all insertions. The number of events analyzed is shown in the tables. ER^+^: estrogen receptor positive tumors. TN: triple negative tumors.

Capture of ectopic sequences at all types of SV junctions is significantly decreased in *BRCA2*-deficient tumors, independently of their estrogen receptor or triple negative status (Fig. 5A and 5B). Moreover, consistent with the data above, the frequency of ectopic sequences capture in INDELs, which do not involve distant DNA, does not decrease in *BRCA2*-deficient cells (Fig. 5A and 5B). These data suggest that in tumors, capture of ectopic sequences associated with SVs (thus involving distant DNA) requires functional BRCA2, although this does not implicate sequence homologies.

The analysis in tumors confirms the implication of BRCA2 in the capture of ectopic sequences at junctions of SVs devoid of sequence homology. This finding fully supports the involvement of the HR players in genetic instability in a sequence homology-independent manner and in a physiopathological context.

## Discussion

HR plays an essential role in genome stability maintenance through processes driven strictly by sequence homology (inspiring the process its name). In contrast to these widely admitted dogmas, the data herein highlight the double face of the pivotal HR actor(s) RAD51 and BRCA2 that intrinsically possess the capacity to foster genetic instability, such as translocations and ectopic sequences capture at SV junctions. In addition, we show here, that 53BP1 protects against RAD51-mediated non-homologous genetic instability. In *Saccharomyces cerevisiae*, HR has been shown to promote genome rearrangements and insertions, but via a process that is strictly dependent on sequence homology (Piazza et al., 2017). Complex chromosomes rearrangements generated by template switch with microhomologies at the junctions were described in yeast, after initiation of BIR and gene conversion, but were found to be independent of RAD51 (Anand et al., 2014; Tsaponina and Haber, 2014). Microhomology-mediated duplications in budding yeast were also found to be independent of HR and EJ machineries (Payen et al., 2008). Here we show that, in human cells, RAD51 can promote both ectopic sequences capture at SVs junctions and SVs themselves, although they do not involve sequence homology, or use only microhomologies far below the MEPS.

We observed these RAD51-mediated non-homologous events in different experimental setups and in the physiopathological context of Brest tumors. Therefore, RAD51 plays contradictory roles: on the one hand, RAD51 maintains genome stability, catalyzing HR but on the other hand, RAD51 can also generate genetic instability in a sequence homology-independent manner. Such genetic instability might account for the poor prognosis associated with RAD51 overexpression in breast, pancreatic, lung, head and neck cancers and colorectal carcinomas (Tennstedt et al., 2013) (and ref in).

Capture of ectopic sequences involves either microhomologies or non-identified patches of nucleotides. Microhomologies-mediated insertions could occur from different processes, i.e. alternative end-joining (A-EJ) or MMTS. Since ablation of HR stimulates A-EJ (So et al. submitted) but decreases microhomologies-mediated insertions (present data), this suggests that microhomologies-mediated insertions occur by MMTS rather than A-EJ.

Because rearrangements and insertions described here involve microhomologies far beyond the MEPS, they cannot be classified as HR events, although they are RAD51-dependent. Instead we propose a novel class of events: non-homologous recombination (nonHR) or micro-homologous recombination (µHR). To reconcile our data with the decades-old knowledge of HR, we propose the model shown in Fig. 6 where nonHR/µHR results from the different attempts of RAD51 to search for homology. Indeed it has been shown *in vitro* that the ssDNA/RAD51 filament scans the duplex DNA for homology through 8-bp blocks (Qi et al., 2015). As in our experiments the median length of microhomologies was from 3 to 6 bp, one can propose that, in living cells, the chromatin context might stabilize, at least transiently, pairing with microhomologies smaller than 8 bp. If a long homologous sequence reaching the MEPS is found, the intermediate is stabilized, triggering canonical HR. Below the MEPS, the intermediate should be rejected, and the ssDNA/RAD51 should engage another round of homology searches. However, as proposed for MMTS (Guirouilh-Barbat et al., 2016), DNA synthesis can be primed by the 3’ invading strand before rejection; therefore, the rejected strand should contain a partial copy of the invaded molecule. Then, this ssDNA can anneal with the other double-strand ends of the recipient molecule, resulting in ectopic sequences capture. Several rounds of homology searches would account for both the observed mosaic sequence capture and the unidentified sequences at the junctions. One can assume that canonical HR should be more stable biochemically and efficient than nonHR/µHR, but this should be largely compensated by the fact that few bp microhomologies are much more frequent than long homologous sequences. In addition, RAD51 increased chromosomal mobility after DNA damage in yeast (Smith et al., 2018). It is thus tempting to speculate that mammalian RAD51 could also do this, favoring inter chromosomal exchange that leads to SVs and capture of ectopic sequences.

**Fig. 6.**
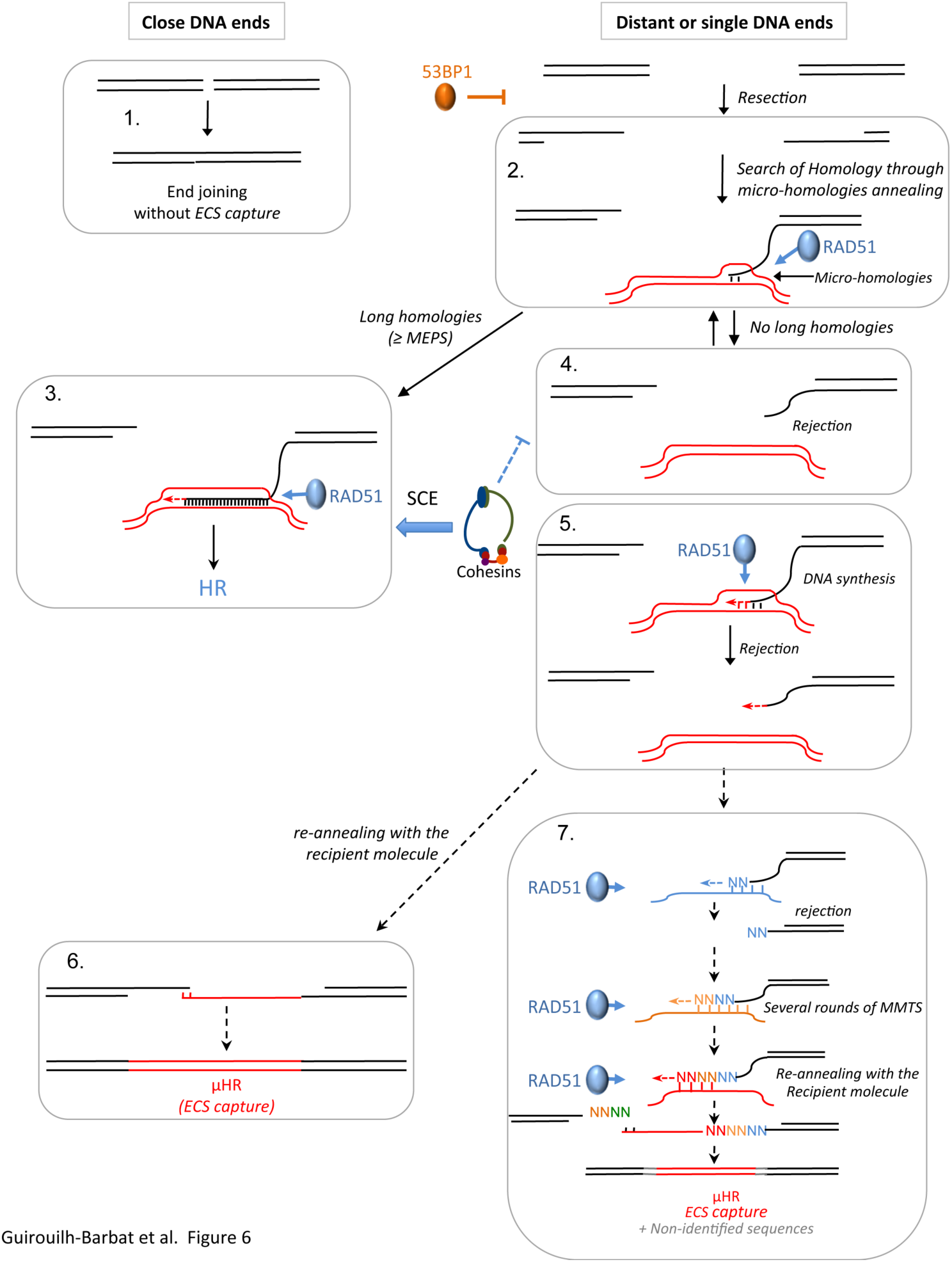
Model for the role of the HR protein RAD51 in non-homologous genetic instability through MMTS. **1/** When two DNA double strand ends are close, insertions (>45bp) were barely recorded (Guirouilh-Barbat et al., 2016). When DNA double-strand ends are distant (few kb), resection generates a 3’ single strand end, which can be covered by RAD51 after loading by BRCA2. One can speculate that similar processes might occur on single-ended DSBs. **2/** The ssDNA/RAD51 filament then searches for homology, starts the scanning for microhomologies, which leads to ectopic matrix invasion. **3/** When a long homology is detected, the intermediate is stabilized, and synthesis occurs for canonical HR repair (left panel). **4/** If no long homology is found, the invading molecule is rejected and re-engaged in another round of homology search (2/). **5/** On some occasions, DNA synthesis can be initiated, as proposed for MMTS. However, the generated intermediate would be very unstable and thus rejected with the copied sequence. **6/** This intermediate re-anneals eventually with the recipient molecule through microhomologies. The resulting product is the insertion of a sequence bordered by microhomologies. **7/** When successive attempts to initiate synthesis by MMTS have occurred, the resulting product contains several sequences (mosaic) and/or the inserted sequences is bordered by unidentified patches of nucleotides. 53BP1 by impairing resection protects against these events. Moreover the cohesion complex represses the capture of ectopic sequences by i) favoring the interaction with the sister chromatids (which bears long identity) which favors sister chromatid exchanges (SCE); ii) by maintaining the two sister chromatids linked limiting the mobility of the broken ends and thus its capacity to interact with an ectopic sequence (Gelot et al., 2016).

One can expect that cells should have developed defense system(s) against RAD51-driven nonHR/µHR. First, we show here that 53BP1 counteracts nonHR/µHR. Recently, Ochs et al. showed that 53BP1 channels DSB repair toward RAD51 dependent HR by preventing hyper-resection (Ochs et al., 2016). Here we show that 53BP1 inhibits microhomology-mediated rearrangements by RAD51, which is an additional way of channeling RAD51 toward faithful HR. However, in breast tumor analysis, all tumors are wild type for 53BP1. This suggests that in a context of high genetic instability such as tumor cells, the protective effect of 53BP1 can be bypassed, at least for the capture of ectopic sequences at SV junctions that involve distant DNA partners. Second, we propose that the cohesion complex also protects against nonHR/µHR. Indeed, by maintaining the sister chromatids’ embrace, the cohesion complex should i) limit the mobility of the DNA ends and thus rearrangements that involve distant sequences; ii) channel the search of homology to the sister chromatid, which shares mega-bases of identity and thus favors sister chromatid exchange by HR. Remarkably, the ablation of the cohesion complex favors the capture of ectopic sequences and genome rearrangements (Gelot et al., 2016). Nevertheless, because nonHR/µHR does not need extended sequence homologies and requires only one DSB (in contrast to rearrangements promoted by end-joining), it might strongly jeopardize genome stability. In addition, because nonHR/µHR can generate mosaic events, it might be involved in complex rearrangements.

In the course of genome analyses, attempts are made to classify the mechanistic origins of the rearrangements in tumors based on the signature of the junctions (Weckselblatt and Rudd, 2015). These classifications should thus be re-evaluated considering the present data.

More generally, as a driver of genome plasticity, RAD51-mediated non-homologous recombination might play important role in and molecular evolution of the genome, in addition to fueling tumorigenesis.

## Methods

### Cells

CG92 and GC49 cell lines were derived from SV40-transformed GM639 human fibroblasts and contained the CD4-3200bp (pCOH-CD4) end-joining reporter (Rass et al., 2009). TERT-RPE-1 cells were cultured in DMEM-F12 (Gibco 10565-018) supplemented with 10% FBS (Gibco 10270106).

### Transfection

The meganuclease I-SceI was expressed by transient transfection of the expression plasmid pCMV-HA-I-SceI (Liang et al., 1998) with Jet-PEI following the manufacturer’s instructions (Polyplus Transfection). The expression of HA-tagged I-SceI was verified by Western blotting. For the silencing experiments, 50,000 cells were seeded 1 day before transfection, which was carried out using INTERFERin following the manufacturer’s instructions (Polyplus Transfection) and 20 nmol siRNA: si53BP1 (onTarget plus SMARTpool for human TP53BP1) was purshased by Dharmacon (Chicago, IL, USA) and Control (5’-AUGAACGUGAAUUGCUCAA -3’), siRAD51#1 (cat# L003530-00-0010, Dharmacon), siRAD51#2 (3’UTR: 5’-GUGCUGCAGCCUAAUGAGA-3’), siBRCA2#1 (5’-GCUGAUCUUCAUGUCAUAA-3’), and siBRCA2#2 (5’-AACUGAGCAAGCCUCAGUCAA -3’) were all synthesized by Eurofins. Forty-eight hours later, the cells were transfected with the pCMV-HA-I-SceI expression plasmid.

### Analysis of DSB repair at the CD4-3200 bp intrachromosomal reporter

After transfection with the pCMV-HA-I-SceI plasmid and incubation for 72 h, the cells were pelleted and resuspended in PBS. After incubation for 1 h at 56°C with 2 mg/ml proteinase K, the proteinase K was inactivated at 95°C for 15 minutes, and genomic DNA was used for the PCR amplification of the junction using the primers CMV-6 (5’-TGGTGATGCGGTTTTGGC-3’) and CD4-int (5’-GCTGCCCCAGAATCTTCCTCT-3’). The predicted size of the PCR product was 732 nt. The PCR products were cloned with a TOPO PCR cloning kit (Invitrogen Life Technologies) and sequenced (Eurofins). For each sample, 2 to 7 experiments were analyzed. In each of these experiments, HA-I-SceI expression and silencing efficiency were verified by Western blot.

### Western blot analysis

Cells were lysed in buffer containing 20 mM Tris-HCl (pH 7.5), 1 mM Na_2_EDTA, 1 mM EGTA, 150 mM NaCl, 1% sodium deoxycholate, 1% (w/v) NP40, 2.5 sodium pyrophosphate, 1 mM β-glycerophosphate, 1 mM NA_3_VO_4_ and 1 μg/ml leupeptin supplemented with a complete mini protease inhibitor (Roche). Proteins (30–40 μg) were denatured, electrophoresed on 9% SDS-PAGE gels, transferred onto nitrocellulose membranes and probed with the following specific antibodies: anti-HA (MMS-101R, Covance), anti-53BP1 (#4937, Cell Signaling), anti-RAD51 (PC130, Millipore), and anti-vinculin (Abcam). BRCA2 depletion was checked on 4-12% Tris acetate gels according to the manufacturer’s instructions and membranes were probed with anti-BRCA2 (#ab9143, Abcam). Immunoreactivity was visualized using an enhanced chemiluminescence detection kit (ECL, Pierce).

### Linear amplification-mediated high-throughput genome-wide translocation sequencing (LAM-HTGTS)

HTGTS was performed according to a published protocol (Hu et al., 2016) and is shown in Figure 2A. Briefly, DNA was purified and sonicated (Bioruptor, Diagenode) into 500–1,000 bp fragments. LAM-PCR was performed using bait primer coupled with biotin, 1U Phusion polymerase (Thermo Scientific) and the following cycles: 1× (98 °C for 120 s); 80× (95 °C for 30 s; 58 °C for 30 s; 72 °C for 90 s); and 1× (72 °C for 120 s). Biotinylated PCR fragments were incubated with MyOne streptavidin C1 beads (Invitrogen) and rotated for 4 h in 1 M NaCl and 5 mM EDTA buffer at room temperature. After washes with B&W buffer (1 M NaCl; 5 mM Tris-HCl, 0.5 mM EDTA), on-bead ligation was performed using 2.5 mM bridge adaptor, 1 mM hexamine cobalt chloride (Sigma), 15 U T4 DNA ligase (Promega), 15% PEG-8000 (Sigma) and the following cycles: 25 °C for 1 h; 22°C for 2 h; and 16°C O/N. After washing three times with B&W buffer (1 M NaCl, 5 mM Tris-HCl (pH 7.4) and 0.5 mM EDTA (pH 8.0)), the on-bead ligated products were subjected to nested PCR using Phusion polymerase, locus-specific and adaptor primers and the following cycles: 1× (95 °C for 300 s); 15× (95 °C for 60 s; 60 °C for 30 s); and 1× (72 °C 600 s). Blocking digestion was then performed with *I-SceI* for 1 h to remove uncut germline DNA. After purification using a Qiagen column, recovered DNA was PCR-amplified using Illumina primers, Phusion polymerase and the following cycles: 1× (95°C for 180 s); 10× (95°C for 30 s; 60°C for 30 s; 72°C for 60 s); and 1× (72°C for 360 s). The tagged PCR products were size-selected for DNA fragments between 500–1000 bp on a 1% agarose gel and purified using a Qiagen column before loading onto an Illumina Miseq machine for paired-end 2X250 bp sequencing.

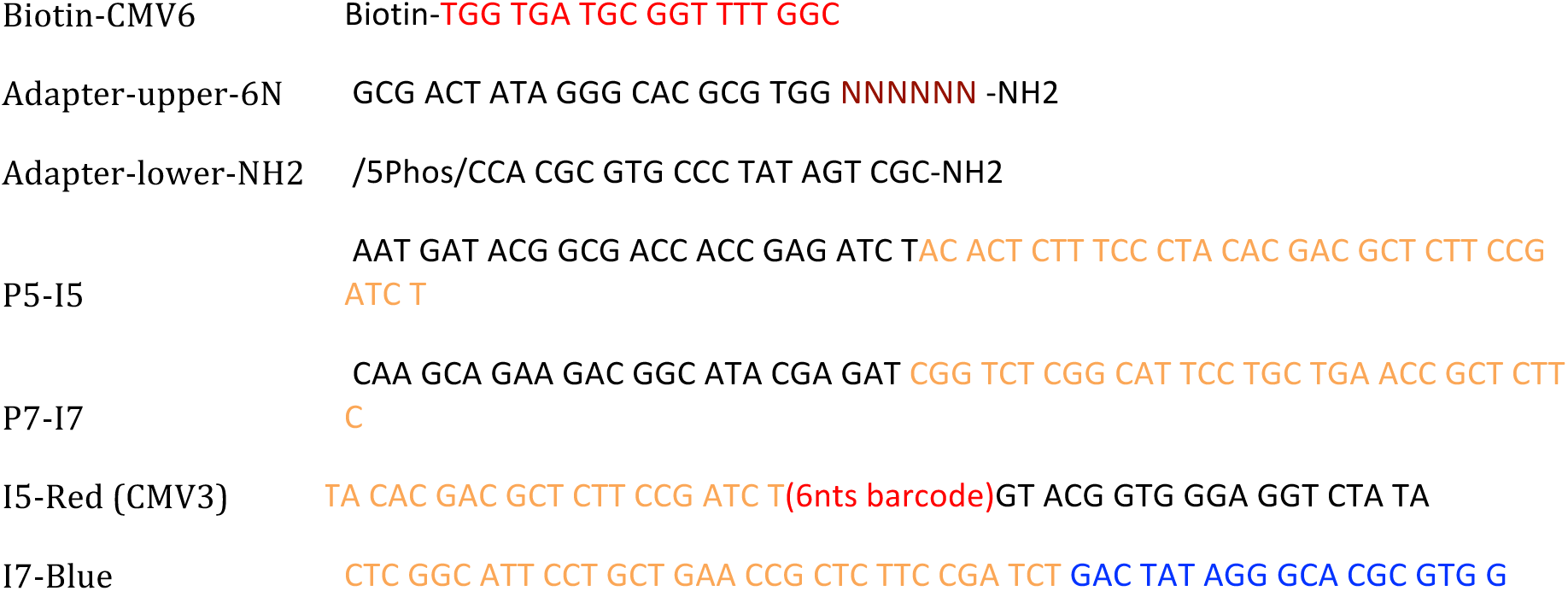
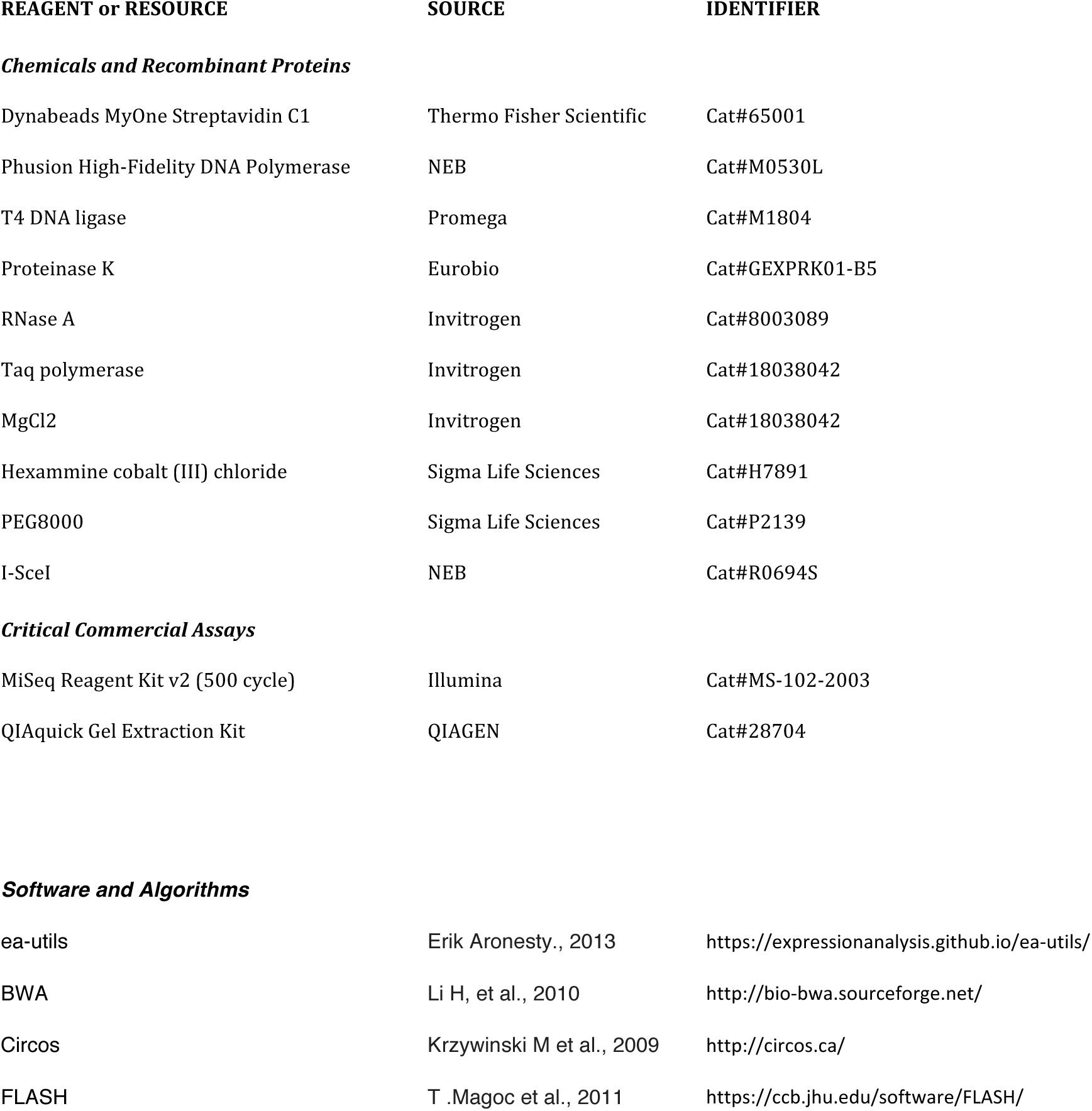

### Data analysis for HTGTS

Sequencing reads in FASTQ files were parallel decoded and adaptor sequence trimmed out by US-utility tool (Aronesty, 2013). Paired-end reads with less than 20-nucleotide bait sequence at the 5’ end of read 1 were filtered out. The remaining paired-end reads were merged by a *FLASH* tool (Magoč and Salzberg, 2011) with a minimum of 10 bp overlap. The merged reads were mapped to the human hg38 reference genome combined with plasmid sequence using BWA (Li and Durbin, 2010) and the default setting. To identify different types of joins, including germlines, deletions, multiple copies and insertions, we first divided the plasmid sequence into several parts, including a blue part, which started with the CMV3 primer and continued until the first I-SceI cut site. The H2Kd part was defined between the first and second I-SceI cut sites. The CD4 part was defined from the second I-SceI cut site to 500 bp downstream, and all rest of the plasmid was defined as the plasmid backbone. According to these definitions, we parsed the mapping of the merged reads. Reads without split mapping and with or without short indels in the first I-SceI cut site were defined as germline. If reads were mapped into two parts in which the 5’ end was blue and CD4 was at the 3’ end, these reads were defined as deletional joins. To identify multiple copies and insertions, the reads must be mapped separately to at least three parts. The reads include the blue part at the 5’ end and the CD4 part at the 3’ end. If one or more pieces of sequence originate from other defined parts but not from the close region (500 bp) of the H2Kd part around the I-SceI cut sites, then these reads are counted as multiple copies and insertions. If the reads included a blue part at the 5’ end and 2 or more pieces of human chromosome sequences, they were also counted as a multiple copy and insertion. All analysis steps above were fulfilled by costumed Perl scripts developed in-house. Circos (Krzywinski et al., 2009) was used to visualize multiple copies and insertions with human chromosome sequences in each 1 kbp bin.

### CRISPR-Cas9 translocation frequency

RPE-1 cells were transfected with 20 pmol siRNA by nucleofection (Amaxa Lonza 4D program EA-104 P3 solution) or Lipofectamine RNAiMAX transfection reagent (ThermoFisher) according to the manufacturer’s instructions. Thirty-six hours after siRNA transfection, 10^6^cells were transfected a second time by nucleofection with pCas9-2A-GFP, sgRNA*^NPM1^* and sgRNA*^ALK^* (3 µg each) (Amaxa Lonza 4D program EA-104 P3 solution). siRNA inhibition was analyzed by Western blot at the time of the second transfection. Transfection efficiency was assessed at 48 h by FACS. First-round PCR was performed on lysates with the primers used for translocation junction detection (listed below). Second-round PCR was performed with SYBR Green and real-time PCR conditions (Mx3005p, Agilent) (Brunet et al., 2009). PCR sequencing was performed on positive wells using forward primers and a BigDye™ Terminator v3.1 Cycle Sequencing Kit according to the manufacturer’s instructions. The sequences were read on an Applied Biosystems™ 3500 Genetic Analyser. At D7, genomic DNA was also extracted from the pool of transfected cells using an E.Z.N.A Tissue DNA Kit (Omega Biotek). Translocation frequencies were estimated by serial dilution as previously described (Renouf et al., 2014).

*sgRNA guide sequences*:

sgRNA*^NPM1^* TATATCCTCGAACTGCTACT

sgRNA*^ALK^* GTCGGTCCATTGCATAGAGG

*Primers sequences*:

*hNPM1-ALK* (PCR1) hNA-Der5-F ACCAAGTCCAGTTGCGTTTC

hNA-Der5-R TTCCCTAACCACTGCCACTC

*hNPM1-ALK* (PCR2) hNA-Der5-NF GAACTCCTGGCCTTAACGTG

hNA-Der5-NR CCACCCTCTAGGGTTGTCAAT

*hALK-NPM1* (PCR1) hAN-Der2-F TCCTTCAGTGTCCATCACGA

hAN-Der2-R GAACCTTGCTACCACCTCCA

*hALK-NPM1* (PCR2) hAN-Der2-NF GAGACATGCCCAGGACAGAT

hAN-Der2-NR AGAGCACATGGGAAAAGGAA

### Whole genome sequence analysis of breast tumors’

We used the data from the Nik-Zainal et al. study (Nik-Zainal et al., 2016) supplementary tables describing 1) clinical annotation 2) structural variation and 3) SNVs and indels found in 560 whole genome sequenced breast cancers. Triple negative, ER-positive HER2-negative high grade ductal carcinoma cases were selected according to clinical annotation. The cases were annotated as HR deficient according to the HRDetect signature (Davies et al., 2017). Cohort of HR proficient cases consisted of 429 cases. The cases were annotated as BRCA2 deficient if mutated in BRCA2 and classified as HR deficient (29 cases, including 19 ER positive and 9 triple negative cases). Non BRCA2 and HR deficient cases were excluded from the analysis. Control group of non-BRCA2 cases consisted of 110 ER positive and 54 triple negative subtypes.

For each category, we analyzed the length of insertions at the SV or INDEL scars.

## Accession number

The raw sequencing data have been uploaded to GEO and the accession number is GSE126546

## Acknowledgments

We are very grateful to Frederick Alt and colleagues for help and advices with setting up the LAM-HTGTS technique This work was supported by grants from the Ligue Nationale contre le cancer “Equipe labellisée 2017”, ANR (Agence Nationale de la Recherche, ANR-14-CE10-0010-02, ANR-16-CE12-0011-02, and ANR-16-CE18-0012-02), AFM-Téléthon and INCa (Institut National du Cancer, PLBIO18-232). L.D. research is supported by the Institut National du Cancer and the Canceropole IdF (#PLBIO16-181), the Institut Pasteur as well as by the European Research Council under the ERC starting grant agreement # 310917.

## Author contributions

Conceptualization: JG-B, BSL; Data curation: WY, TP; Funding acquisition: BSL, LD; Investigation: JG-B, WY, LB, EYG, AS, CL, EB; Methodology: JG-B, YW, TP, CL, MHS, EB, LD, BSL; Project administration: BSL; Resources: MHS, EB, LD, BSL; Supervision: JG-B, TP, EB, LD, BSL; Validation: JG-B, BSL; Visualization: JG-B;, WY, EB, LD, BSL; Writing – original draft: JG-B, BSL; Writing – review & editing: JG-B, WY, TP, CL, EB, LD, BSL.

## Supplemental data S1

**Fig. S1. - Sequence of the junction scars at the I-SceI site of the CD4-3200bp substrates**.

**S1A. Sequences of captured ECS at junction scars of the two I-SceI sites in GC92 cells bearing the CD4-3200bp reporter after 53BP1, RAD51, and/or BRCA2 siRNA transfection**. Insertions were blasted using the BLAST program of the National Centre of Biotechnology Information, National Institutes of Health, Bethesda MD, USA). Insertions were blasted to the End Joining reporter, the I-SceI expression plasmid, the mitochondrion genome, the human genome (*Homo sapiens*, taxid: 9606), Human ALU repeat elements and the Nucleotide collection, using megablast and discontinuous megablast. Sequences identified as “unmappable stretches of nucleotides” were identified by neither of these searches.

In red : microhomologies ; in yellow: unmappable stretches of nucleotides ; lower case : sequence of the CD4-3200bp reporter.

S1A. Sequences of insertions

in red : microhomologies

in yellow: non template

lower case : sequence of the CD4-3200bp reporter

siCTRL

155nt

1-154: Homo sapiens chromosome 16, alternate assembly CHM1_1.1 3221396 to 3221549

tttggcCCAACAGGATTTGACCCTGAGGCCCACTCTCACCCTAATCATAACCGCAAAACCACCAGCGCCTGGAGAGAGAGTGAGAGAGAAACAGAAACGGAGCGAGTGTGTGTCGCTGTGATGCCCTTTGCCGTCGCTGCTCATCCCCAGTGACCTCCTGA

395 nt

1-395: Sequence répétée non Alu Homo sapiens chromosome 12, alternate assembly CHM1_1.1 48768696 to 48769089

tggtgatgcggttttggcGTGGATGATCTTTTTGTTGACGTTGATGCTATTCCTGTTTGTTGGTTTTCCTTCTAACAGTCAGGCCCCTCAGCTGCAGGTCTGTTGGAGTTTGCTGGAAGTCCACTCCAGACTCTACTTGCCTGGGTATCACCAACGGAGGCTGCAGAACACCAAATATTGCTGCCTGATCCTTC CTTCCTCTAGAAGCTTCATCACAGAGGGGCACCCACCTGTTTGAGGTGTCTGTCGGCCCCTACTGGGAGGTGTCTCCCAGTCAGGCTACACAGGGGTCAAGGACCCGCTTGAGGAGGCAGTCTGTCCATTTTCGGATCTTGAACGCCATGCTGAGAAAACCACTGCTCTCTTCAGAGCTGTCAGATAGGGACGTTTTAAGTCTGCAGAAGCTGTCTGCTgc

117nt

3-70 CD4-3200bp reporter ; 71-117 CD4-3200bp reporter (different location)

TATCGGCCTCTGAGCTATTCCAGAAGTAGTGAGGAGGCTTTTTTGGAGGCCTAGGCTTTTGCAAAAAGCTAACTTGTTTATTGCAGCTTATAATGGTTACAAATAAAGCAATAGCATtatccct

298 nt

16-290: Homo sapiens chromosome 3, alternate assembly CHM1_1.1 12139590 to 12139864

TCACTGCCACCAGAATCAAGCGATTCTCCCGCCTCAGCCTCCTGAATAGCTGGGATTACAGGCGCCTGCCACCATGCTTGGCTAATTTTTTGTATTTTTAGTAGAGACAGCGTTTCACTATGTTGACCAGGCTGGTCTTGAACGCCTGACCTCGTGATCCACCTGCCTCAGCCTCCCAAAGTGCTGGGATTACAGGCGTGAGCCACCACGTCTGGCCACAAATTTCTTTAAGAGAACCCATATACTAGGAGCCTGTAACAACCAGGCTGTCAAAGTGGGGCAGCGGTGCTGATGTTAG

220nt

1-223: Homo sapiens chromosome 7, alternate assembly CHM1_1.1 72739670 to 72739855 ; 1-186:Alu sequence

tggtgatgcggttttggcTCACTGCAACCTCCGTCTCCCAGGTTCAAGCGATTCTCATGCCTCAGACTCCCGAGTAGCTGAGATTACAGGCGTGCGCCACCATGCCTGGCTAATTTTTGTATTTTTAGTAGAGACGGTGATTCACCATGTTGGCCAGGCTGGTCTTGAACTCCTGACCTCAGATGATCTGCCAACCTCGCCCTCATAGGCCTGAGATTTTAAAGCATGCGTGGGAATATATAGTTTAG

164nt

1-160: Homo sapiens chromosome 16, alternate assembly CHM1_1.1 3221396 to 3221556

tttggcCCAACAGGATTTGACCCTGAGGCCCACTCTCACCCTAATCATAACCGCAAAACCACCAGCGCCTGGAGAGAGAGTGAGAGAGAAACAGAAACGGAGCGAGTGTGTGTCGCTGTGATGCCCTTTGCCGTCGCTGCTCATCCCCAGTGACCTCCTGAACTTTCAGG

154nt

1-153: Homo sapiens chromosome 16, alternate assembly CHM1_1.1 3221396 to 3221548

tttggcCCAACAGGATTTGACCCTGAGGCCCACTCTCACCCTAATCATAACCGCAAAACCACCAGCGCCTGGAGAGAGAGTGAGAGAGAAACAGAAACGGAGCGAGTGTGTGTCGCTGTGATGCCCTTTGCCGTCGCTGCTCATCCCCAGTGACCTCCTA

317 nt

4-317 CD4-3200bp reporter

AACTGCTGATGCCCGTTTGGCAGTACATCAATGGGCGTGGATAGCGGTTTGACTCACGGGGATTTTCAAGTCTCCACCCCATAGACGTCAATGGGAATTTGTTTTGGCACCAAAATCAACGGGACTTTCCAAAATCTAGTAACAACTCCGCCCCATTGACCCAAATGGGCGGTAGGCGTGTACGGTGGGAGGTCTATATAAGCAGAGCTCTTTGGCTAACTAGAGAACCCACTGCTTACTGGCTTATCGAAATTAATACGACTCACTATAGGGAGACCCAAGCTGGCTAGCGCTCTAGAGCAACACGGAAGGAATTAatgtgccgag

157 nt

1-157: CD4-3200bp reporter

ggaaggaCAGAAAGCGGGTCAACGGTGAGGCCATGGTGTGCTCTCCCCTAGAGCCCTACGGAATTCGAGCTCGCCCGGGGATCTCGAGGTCACCCTGCGGTGTCGTCCATCACAGTTTGCCAGTGATACACATGGGGATCAGCAATCGCGCATAATCGATATTAccctatcc

113 nt

1-110: Homo sapiens chromosome 8, GRCh38.p12 Primary Assembly 136408078 to 136408187

ggaattCCTAAGAGCAGAACTATAGTTTGTCACCTCCACAACTCCAGGGACTGGCACTAGATTGGCATATTTTTCAGTAAATGTTGAATAAAGGAATGAATCTTACTTCAAAGAGGTCC

107 nt

1-99: seq répétée

ctgttatGAAGGCCACAAAGTGGTCCAAATATCCACTTGCAGATTCTACAAAAAGAGTGTTTGAAAGCTGAACTATGAAAGCAAGGTTCAACTCTGTGAGTAATTCCTTCCGTG

si53BP1

130nt

1-61: CD4-3200bp reporter; 62-129: CD4-3200bp reporter (different location)

cctgttatTTGTGAAATTTGTGATGCTATTGCTTTATTTGTAACCATTATAAGCTGCAATAAACAAGTTAGCTTTTTGCAAAAGCCTAGGCCTCCAAAAAAGCCTCCTCACTACTTCTGGAATAGCTCAGAGGCCGATccctatctag

130nt

1-61: CD4-3200bp reporter ; 62-129: CD4-3200bp reporter (different location)

ccctgttatTTGTGAAATTTGTGATGCTATTGCTTTATTTGTAACCATTATAAGCTGCAATAAACAAGTTAGCTTTTTGCAAAAGCCTAGGCCTCCAAAAAAGCCTCCTCACTACTTCTGGAATAGCTCAGAGGCCGATccctatctag

130nt

1-61: CD4-3200bp reporter ; 62-129: CD4-3200bp reporter (different location)

ctgttatTTGTGAAATTTGTGATGCTATTGCTTTATTTGTAACCATTATAAGCTGCAATAAACAAGTTAGCTTTTTGCAAAAGCCTAGGCCTCCAAAAAAGCCTCCTCACTACTTCTGGAATAGCTCAGAGGCCGATccctatctag

254nt

1-222: Homo sapiens chromosome 8, alternate assembly CHM1_1.1 94640757 to 94640978

cacggaaggGAAGAGGGAAAGGACATGGAATGTTTGAGAAGCAAACACACACTTCTCCCCAGAGTGATTCTGTGCATTAACCAGACACGTCTCCATTCTGCTCCCATGGACCTACTGCTGGTGATTCCCATTTGCAGTCGCCTAACCTACTACCCTTACGCTGTTCTCTGCTGTGTTCCTCAACCATCAGCATTTGGGGCAATTCTTCACAGCACAGGACTGCCCTCATATTTTAAATAAGAAATGTGTGATAACATCAAAAG

154nt

1-153: Homo sapiens chromosome 16, alternate assembly CHM1_1.1 3221396 to 3221549

tttggcCCAACAGGATTTGACCCTGAGGCCCACTCTCACCCTAATCATAACCGCAAAACCACAGCGCCTGGAGAGAGAGTGAGAGAGAAACAGAAACGGAGCGAGTGTGTGTCGCTGTGATGCCCTTTGCCGTCGCTGCTCATCCCCAGTGACCTCCTGA

237 nt

1-140 Homo sapiens chromosome 19, alternate assembly CHM1_1.1 5640605 to 5640746 ; 1-41: Alu sequence; 40-140: Alu sequence; 140-200: Alu sequence ; 40-215: Homo sapiens chromosome 19, alternate assembly CHM1_1.1 1832221 to 1832391

tttggcTCACTGCAACCTCCACCTCCTGGGTTCAAGCGATTCTCCTGTATTTTTAGTAGAGATGGGGTTTCACCATGCTGGCCAGGCTGGTCTTGAACTCCTGACCTGGTGATCTGCCCGCCTCGGCCTCCCAAAGTGCTGGGATTGCCTCAGCCTCCCAAGCAGCTGAGATTACAGGCATGCACCACCATGCTCGGCTAATTTTTACAGGCGTGAGCCACTGCACCCGGACCACACCTAGCA

238 nt

1-224 Alu sequence

tttggcTCACTGCAACCTCCACCTCCTGGGTTCAAGCGATTCTCCTGCCTCAGCCTCCCAAGCAGCTGAGATTACAGGCATGCACCACCATGCTCGGCTAATTTTTGTATTTTTAGTAGAGATGGGGTTTCACCATGCTGGCCAGGCTGGTCTTGAACTCCTGACCTGGTGATCTGCCCGCCTCGGCCTCCCAAAGTGCTGGGATTACAGGCGTGAGCCACTGCACCCGGACCACACCTATGCA

241nt

1-224 Alu sequence

tttggcTCACTGCAACCTCCACCTCCTGGGTTCAAGCGATTCTCCTGCCTCAGCCTCCCAAGCAGCTGAGATTACAGGCATGCACCACCATGCTCGGCTAATTTTTGTATTTTTAGTAGAGATGGGGTTTCACCATGTTGGCCAGGCTGGTCTTGAACTCCTGACCTGGTGATCTGCCCGCCTCGGCCTCCCAAAGTGCTGGGATTACAGGCGTGAGCCACTGCACCCGGACCACACCTATGCATCG

386nt

1-384 Homo sapiens chromosome 8 alternate assembly CHM1_1.1 29167094 to 29167478

gcggttttggcTCACTGCAGGCTCCGCCCCCCGGGGTTCACACCATTCTCCTGCCTTAGCCTCCCGAGTAGCTGGGACTACAGGTGCCCGCCACCTCGCCTGACTAGTTTTTTGTATTTTTAGTAGAGACGGGGTTTCACCGTGTTAGCCAGGATGGTCTCGATCTCCTGACCTCGTGATCCGCCCTCCTTGGCCTCCCAAAGTGCTGGGATTACATACGTGAGCCACTGCGCCCAGCCTGAAGTTTCTTTTCTGA CAGAAATAGGAAGGAAATTTAGACAGAAGTACGAGCTTTAGCAGAAAAATTGTTTTTCAACCCCAGTAGTCAGAGTCTAATAAACCTTATTTTAATTGGCAAATTAAAAGAAATGAACTCTAAAACACTTCAAGTAAGTCA

69nt

1-56: Homo sapiens chromosome 8 GRCh38.p1285269931 to 85269986

gtcgtaTTATATTGTAACCCACAATATCGTTTACAGGCAAAATAATGGTATATCATGAGACAGATTGTTTGGCCG

185nt

1-184: Alu sequence

gttttggcTCACTGCAACCTCCACCTCCCAGGTTCAACAATTCTCGTGTCTCAGCCTCCCAAGTAGCTGGGACTACAGGCGCCTGCCGCCACACCTGACTAATTTTTGTATTTTTGGTAGAGACGGAGTTTCCCTATGTTGCCCAATCCGGTCTCGAACTCCTGGGCTCAAGTGATGAGCCCACCTCGGCCT A

94nt

33-94 : Homo sapiens chromosome 1, GRCh38.p12 Primary Assembly 86576654 to 86576715

ggtctatataCACAGAGCTCTCGGCTCTTGGTGATGCGGTTTTGGCAGGAAAAGAAAGCAGGAGTAATCCTCGACAGCTGGAACAAAGTATAAATATAGTCCTGagaactg

114 nt

1-108 repeated sequence

ATTGATTATTCCATTACATTCCATTCGATGATTCCATTTGAGTCCATTCAATGATTCTATTCGATTCCATTCAATAATTCCATTCGATTCCATTTGATGATGATCCATCTCACG

167 nt

1-159: Homo sapiens chromosome 16, GRCh38.p12 Primary Assembly 3171420 to 3171578

gatgcggttttggcCCAACAGGATTTGACCCTGAGGCCCACTCTCACCCTAATCATAACCGCAAAACCACCAGCGCCTGGAGAGAGAGTGAGAGAGAAACAGAAACGGAGCGAGTGTGTGTCGCTGTGATGCCCTTTGCCGTCGCTGCTCATCCCCAGTGACCTCCTGAACTTCAGAGCTCc

266 nt

1-99: : Homo sapiens chromosome 16, GRCh38.p12 Primary Assembly 3681336 to 3681434 ; 96-255: Homo sapiens chromosome 16, GRCh38.p12 Primary Assembly

ggtgatgcggttttggcaCCCATTCTGCCTGTGCTGCCTACAATGGGAAGGCTGACAGCTGGCCAGTCTCCCTGGCCTGCTGCTTGCTGTGGATTATTTGCTGAGTGGCACTGCCCATTCTGCCTGTGCTGCCTACAATGGGAAGGCTGACAGCTGGCCAGTCTCCCTGGCCTGCTGCTTGCTGTGGATTATTTGCTGAGTGGCACTGACCACCCTTGCTTTGCCAATGGAGACAGCACTGGCGCCTGGTGCTAGTGGGCTCCCAAGGCTCCTTTCCGGGTTCT

197 nt

1-180: CD4-3200bp reporter

ggagacccaagctgTGGTGATACGGTTTTGGCAGTACATCAATGGGCGTGGATAGCGGTTTGACTCACGGGGATTTCCAAGTCTCCACCCCATTGACGTCAATGGGAATTTGTTTTGGCACCAAAATCAACGGGACTTTCCAAAATGTCGTAACAACTCCGCCCCATTGACGCAAATGGGCGGTAGGCGTGTACTGGCTTATCGAAAACAC

99 nt

1-99 : pCOH 45084606

gaaggaattCGAGCTCGCCCGGGGATCTCGAGGTCACCCTGACGGTGTCGTCCATCACAGTTTGCCAGTGATACACATGGGGATCAGCAATCGCGCATAATCGATATTaccctgtta

99 nt

1-99 : pCOH 45084606

gaaggaattCGAGCTCGCCCGGGGATCTCGAGGTCACCCTGACGGTGTCGTCCATCACAGTTTGCCAGTGATACACATGGGGATCAGCAATCGCGCATAATCGATATTaccctgttat

354 nt

1-354 : Homo sapiens chromosome 14, GRCh38.p12 Primary Assembly 75197885 to 75198238, 1-95 repeated sequence

ggttttggcaTGTTGGCCAGACTGGTCTCGAATTCCTGACCTCAGGTGATCCGCCCGCCTCGGCCTCCCAAAATGCTGGAATTATAGGCGTCAGCCACTGCGCCCTGCTAATGGACAACCTTTAAACCTACATGTTTGGCTATTTCTTAGTATTTTAGGGTTGCTTCGTGCTCTGGCCTGGGTTTGCAGGAAAAATGGGAGAGAAGAGGAGAAAAGGGCTGGAAGAGGCTTGGCCATGGTAGAAGGCATGGGTG GGTGTGAGGAGAAGGAAACAGAGTGATGGGCCAGCAAGGAGTCACAGGGTTTCGGGATCTGGGCTAGCGACCCTCCTGGCCTCCAATGGGCTAACACGGAAAAGAGGGATc

129 nt

1-129: Homo sapiens chromosome 9, GRCh38.p12 Primary Assembly 99847500 to 99847628

ggttttggcaAGGTGGTGCTTACCTGTGATCATGCTCAGTGCTGACAGGCAGGCTAAGGCTTGGATATCAAGGTTCAGGCTCTGCAAATTTAAGGAAAAGTCTTTAATAGAGTCGAACCACTCCCCAAATCCACGAAGG

siRAD51 #1

none

siRAD51#2

49nt

1-49 pBACSe

ccctgttCACACAGGAAACAGCTATGACCATGATTACGCCAAGCTTGGGCTGCAGGtccctatc

1-130 Homo sapiens chromosome 5, GRCh38.p12 Primary Assembly 172716876 to 172717005

ttaccctCTTGGGAGCCGACAAGGGGGTCTCTAGCGCCACCTTCCGGGCATCCGGGAAATGTCCGTACAGTCCCACCAAGGCCCACCTGCCGCCTCGACATTCATAAATAAACGCCAACAATGACAGCGACAGCAACTGGTTATTACCCTCAG

1-140 pCOH CD4 190-329

cctgttatcGTGCACCCAACTGATCTTCAGCATCTTTTACTTTCACCAGCGTTTCTGGGTGAGCAAAAACAGGAAGGCAAAATGCCGCAAAAAAGGGAATAAGGGCGACACGGAAATGTTGAATACTCATACTCTTCCTTTTTCAATATC

177 nt

72-123: Homo sapiens chromosome 9, GRCh38.p12 Primary Assembly 16273586 to 16273637

GAAAGAGTTCATTGTTTCTGATTGTAAAAATAAAACTTAATATATATGCTCTTCCAAGAAGAATCACTGACTACATAAATGGATAAAGAAGAAAAGGCAATTCATCCATAGTCCTGCCACACAGAGAAAGCCATTCTTCGTATTTTGGCATATGTACTTTTTAATAAAAGTATACTT

156 nt

1-154 : Homo sapiens chromosome 16, GRCh38.p12 Primary Assembly 3681281 to 3681434

ttttggcaCCCATTCTGCCTGTGCTGCCTACAATGGGAAGGCTGACAGCTGGCCAGTCTCCCTGGCCTGCTGCTTGCTGTGGATTATTTGCTGAGTGGCACTGACCACCCTTGCTTTGCCAATGGAGACAGCACTGGCGCCTGGTGCTAGTGGGCTCCCATGCT

si53BP1+ siRAD51#1

none

si53BP1+ siRAD51#2

214 nt

1-210: CD4-3200bp reporter

cgctctagagcaacTGATCTTCAGCATCTTTTACTTTCACCAGCGTTTCTGGGTGAGCAAAAACAGGAAGGCAAAATGCCGCAAAAAAGGGAATAAGGGCGACACGGAAATGTTGAATACTCATACTCTTCCTTTTTCAATATTATTGAAGCATTTATCAGGGTTATTGTCTCATGAGCGGATACATATTTGAATGTATTTAGAAAAATAAA CAAATAGGGGTTGGAAcctatctagatat

48 nt

1-48: CD4-3200bp reporter

cgactcactatagATATGAAATCACGCCATGTAGTGTATTGACCGATTCCTTGCGGTCCGAtctagatatgaa

39 nt

39 nt : non template

accctgttatATTATATCCCTATCTAGGAATTACCCTGTTATATTATATccctatctag

124 nt

1-124: pBSCe ISceI

accctgttaCCATCATCCATGAACCAGTATGCCAGAGACATCGGGGTCAGGTAGTTTTCAACCAGGTTGTTCGGGATGGTTTTTTTGTTGTTAACGATGAACAGGTTAGCCAGTTTGTTGAAAGCTTGGTGTTtccctatc

siBRCA2 #1

153 nt

1-154 Homo sapiens chromosome 16, alternate assembly CHM1_1.1 3221396 to 3221549

atgcggttttggcCCAACAGGATTTGACCCTGAGGCCCACTCTCACCCTAATCATAACCGCAAAACCACAGCGCCTGGAGAGAGAGTGAGAGAGAAACAGAAACGGAGCGAGTGTGTGTCGCTGTGATGCCCTTTGCCGTCGCTGCTCATCCCCAGTGACCTCCTGAA

siBRCA2 #2

111 nt

9-111: pBASCe

GACGGCTTGTTTCTTTTCTGTGGCTGCGTGAAAGCCTTGAGGGGCTCCGGGAGGGCCCTTTGTGCGGGGGGAGCGGCTCGGGGGGTGCGTGCGTGTGTGTGTGCGTGGGGAtatctagat

164 nt

3-164: pCOH 1362-1201

CACGTGTTGCTCTAGAGCGCTAGCCAGCTTGGGTCTCCCTATAGTGAGTCGTATTAATTTCGATAAGCCAGTAAGCAGTGGGTTCTCTAGTTAGCCAGAGAGCTCTGCTTATATAGACCTCCCACCGTACACGCCTACCGCCCATTTGCGTCAATGGGGCGGAG ttcgaattcgag

si53BP1+ siBRCA2 #1

132 nt

17-132: Bovine satellite DNA fragment

TATTGATCACGTGGCTGATCATGCACTGATCACGTGACTATCATGCACTGCTCACGTGGCTGATCATGCGATGATCACGCAGCTACCATGTACTGGTCACATGATTGATCATGAACTGATCACGTGACTGATtatccctatc

120 nt

1-120: Homo sapiens chromosome 1, alternate assembly CHM1_1.1 86165308 to 86165427

acggaaggaattTCCCAATACCGTTCCTAAAAATTAGAACTACTTAGAAATTTACCCTTGGCTGTTATGAGGAAAAATCTGAGTCTATAGGAGTCAGTTTGTTTTCACTAAATGAGCTTGAAGCATGACTTGtgttatcccta

66 nt

4-67: pCOH 1300-1363

AGATATCGAAATTAATACGACTCACTATAGGGAGACCCAAGCTGGCTAGCGCTCTAGAGCAACACGGtcacccattcgaatt

si53BP1 + siBRCA2 #2

91 nt

6-85: Homo sapiens chromosome 5, alternate assembly CHM1_1.1 35891395 to 35891481

CTTGGTATGTATGTATGTATGCATACATATGTATGCATACATGTATACATATGGTATACATATGGTATGCATGTATGTAGGCATACCAATA

156 nt

1-154: Homo sapiens chromosome 16, alternate assembly CHM1_1.1 3221396 to 3221549

ggttttggcCCAACAGGATTTGACCCTGAGGCCCACTCTCACCCTAATCATAACCGCAAAACCACCAGCGCCTGGAGAGAGAGTGAGAGAGAAACAGAAACGGAGCGAGTGTGTGTCGCTGTGATGCCCTTTGCCGTCGCTGCTCATCCCCAGTGACCTCCTGAA

**S1B. Sequences of all junctions of the two I-SceI sites in GC92 cells bearing the CD4-3200bp reporter after 53BP1, RAD51, and/or BRCA2 siRNA transfection (see excel document).**

**Fig. S2.**
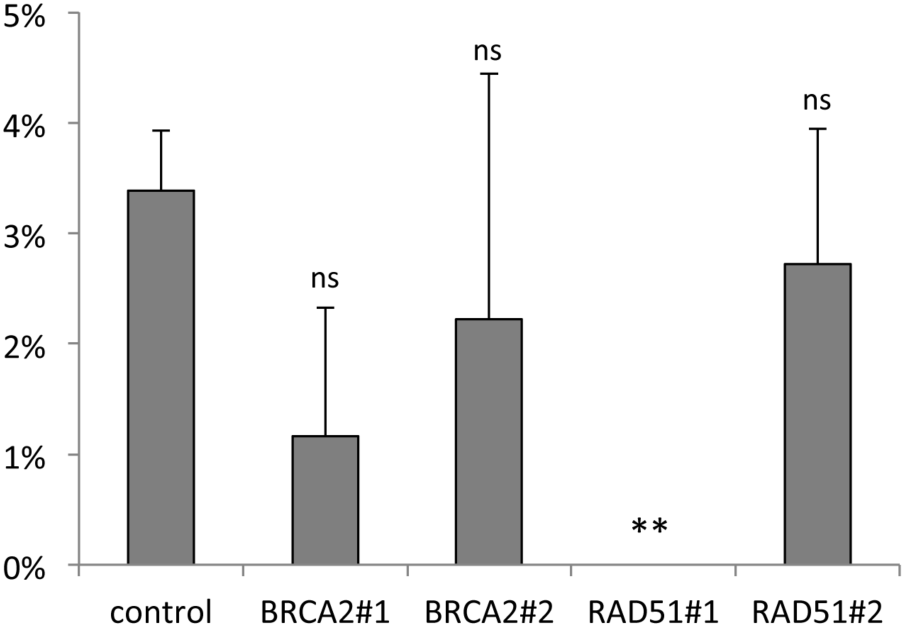
ECS capture in the presence of 53BP1 after BRCA2 or RAD51 depletion in GC92 cells. Bars represent the average of 2-8 experiments (mean ± SEM).

**Figure S3.**
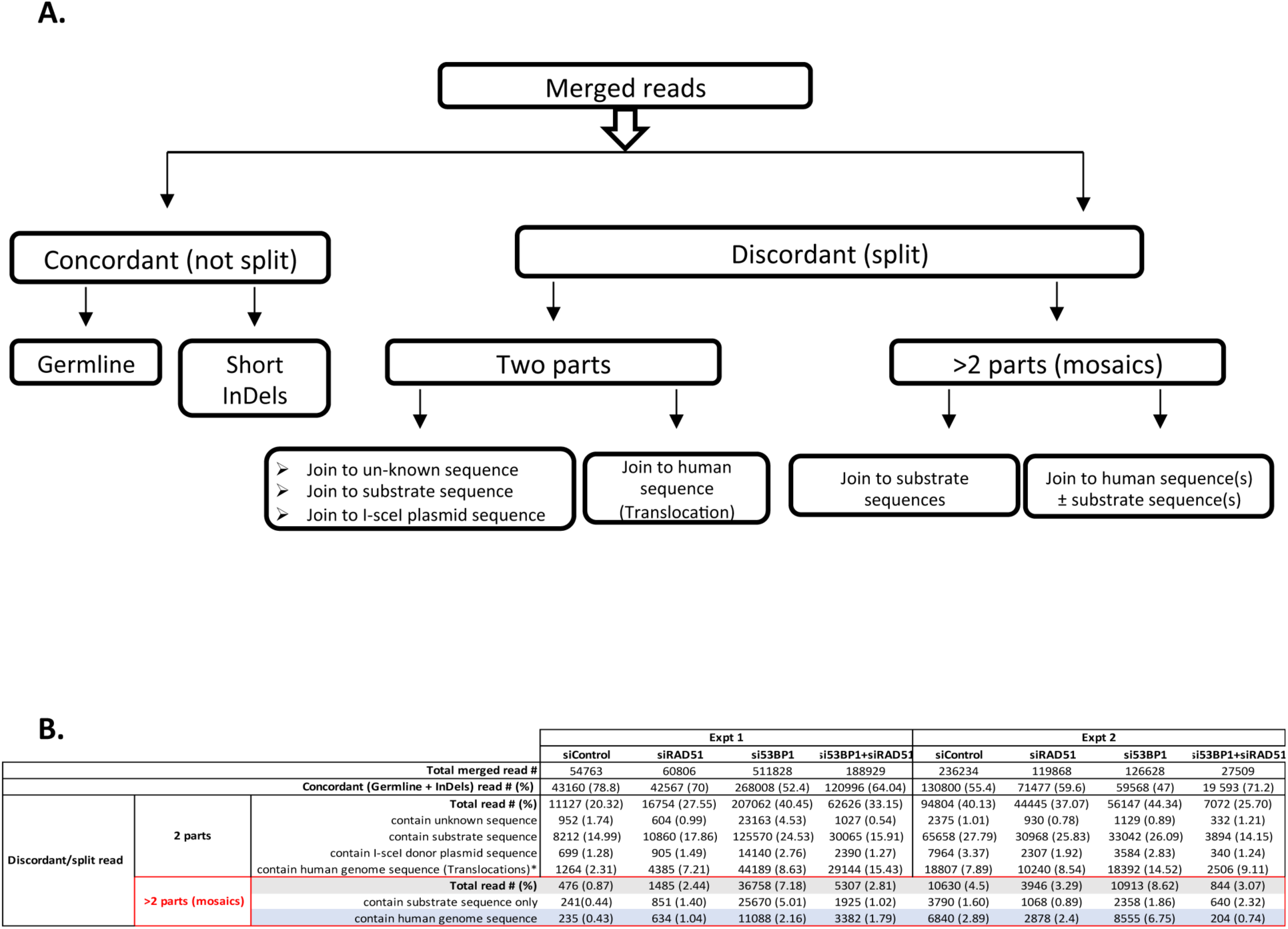
A. Scheme of LAM-HTGTS analysis pipeline. B. Numbers of LAM-HTGTS-sequenced and analyzed reads.

**Fig. S4.**
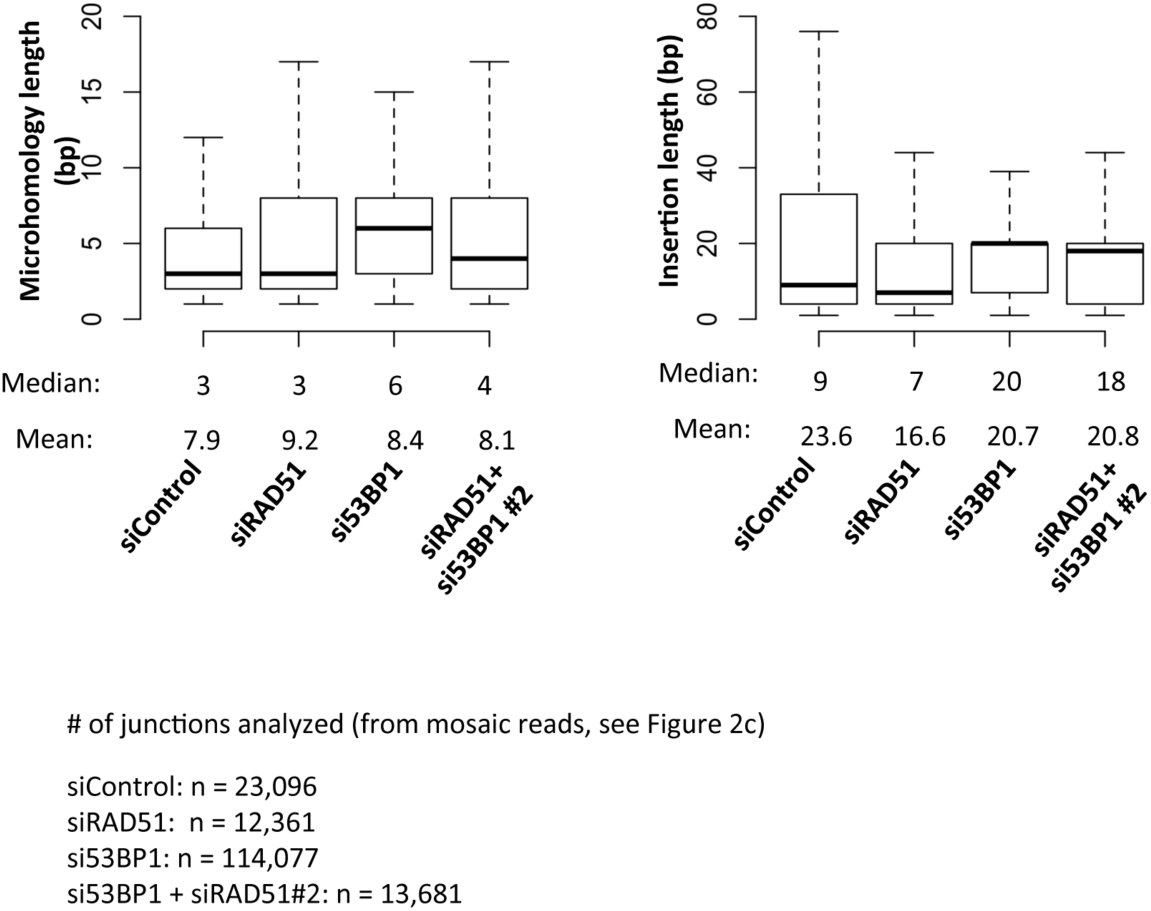
Microhomology (left panel) and insertion length (right panel) in the junction sequences of mosaic reads. The thick horizontal line indicates the median, and the upper and lower lines correspond to the 25th and 75th percentiles.

**Fig. S5.**
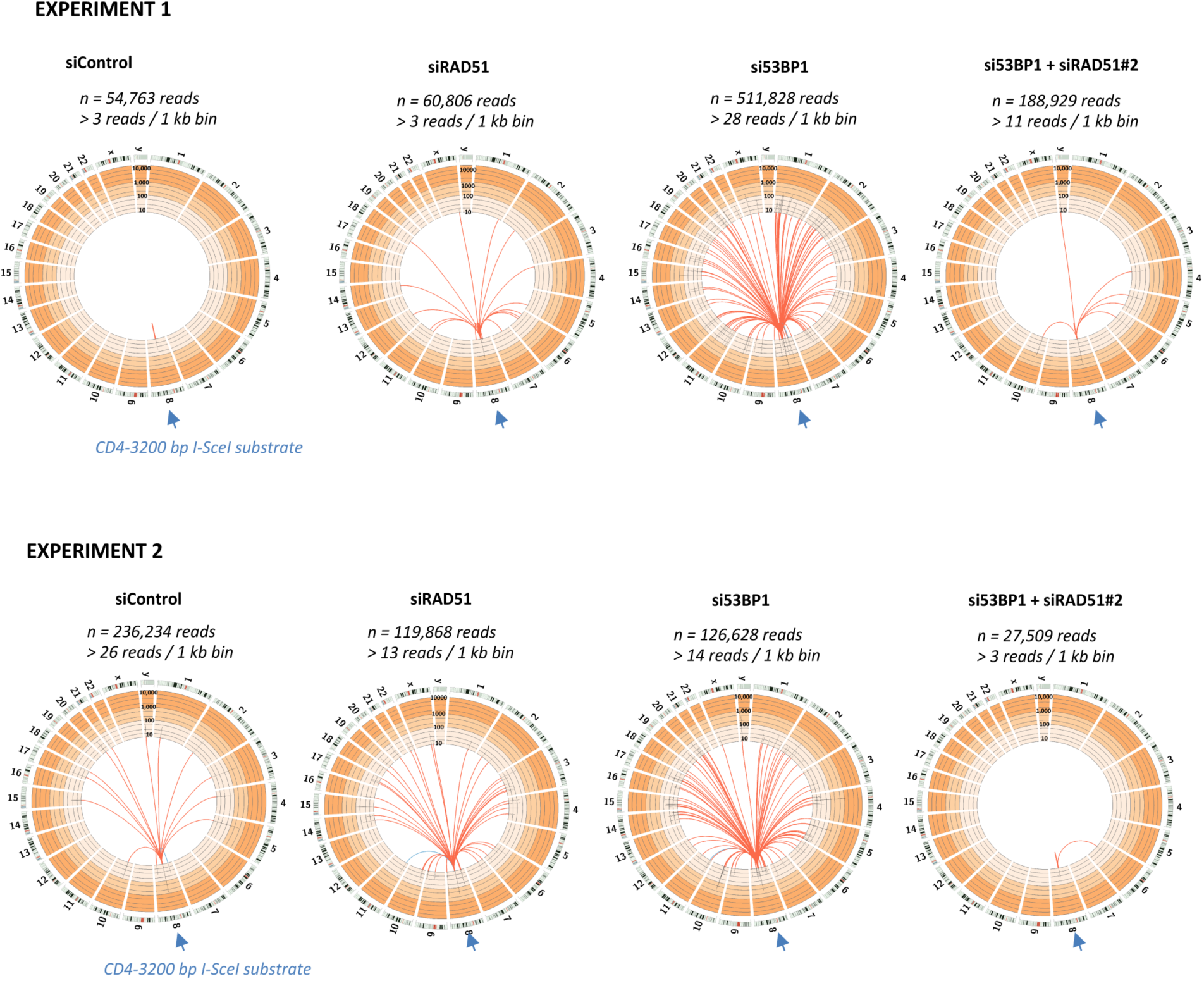
Biological duplicates of the LAM-HTGTS experiments. Circos plots represent interchromosomal junctions.

**Fig. S6.**
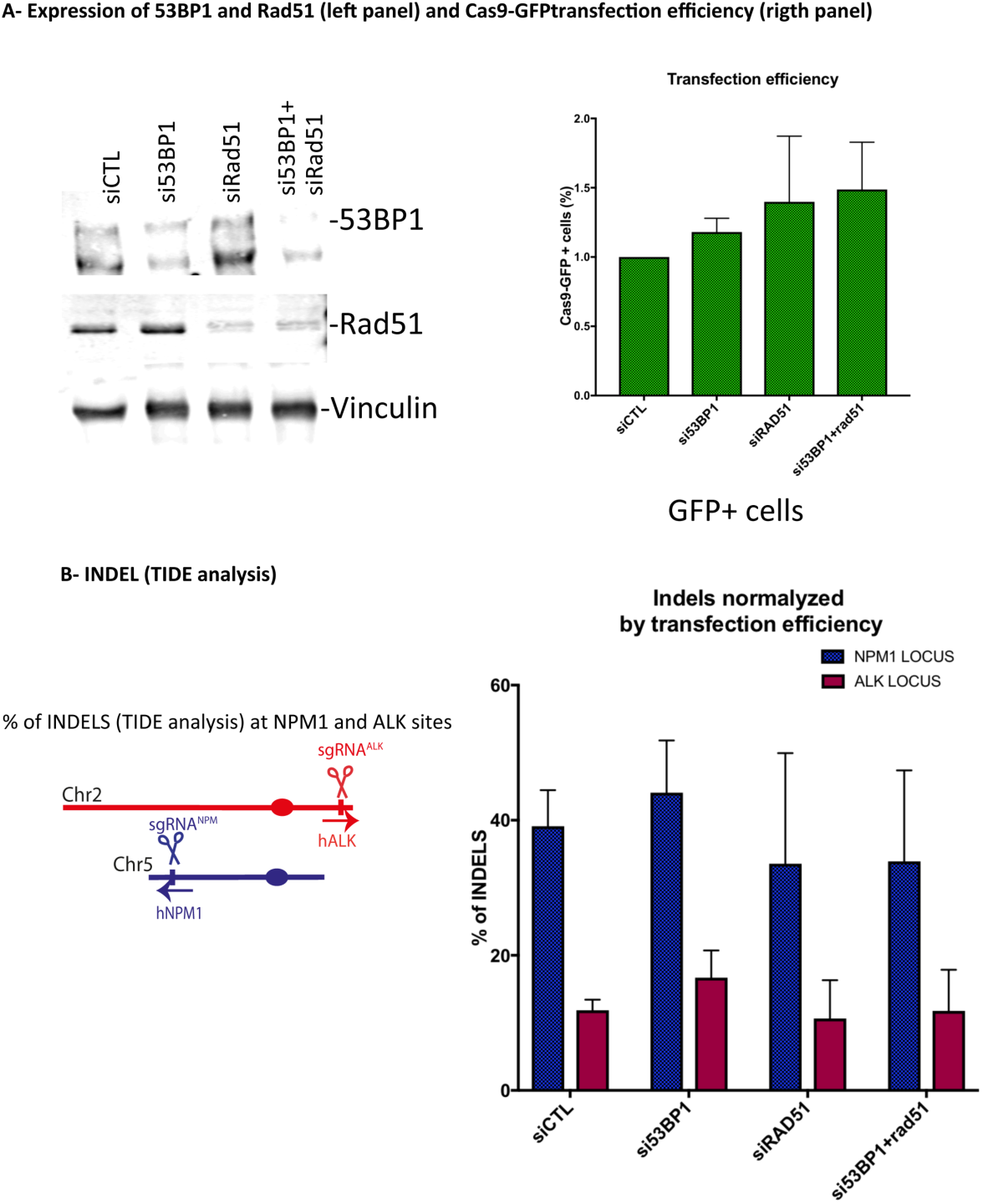
A) Analysis of 53BP1 and Rad51 expression (A-left panel) and Cas9-GFP transfection efficiency (A-right panel) and (B) % of INDELs corresponding to CRISPR /Cas9 cleavage efficiency upon silencing of RAD51 and/or 53BP1.

**Fig. S7.**
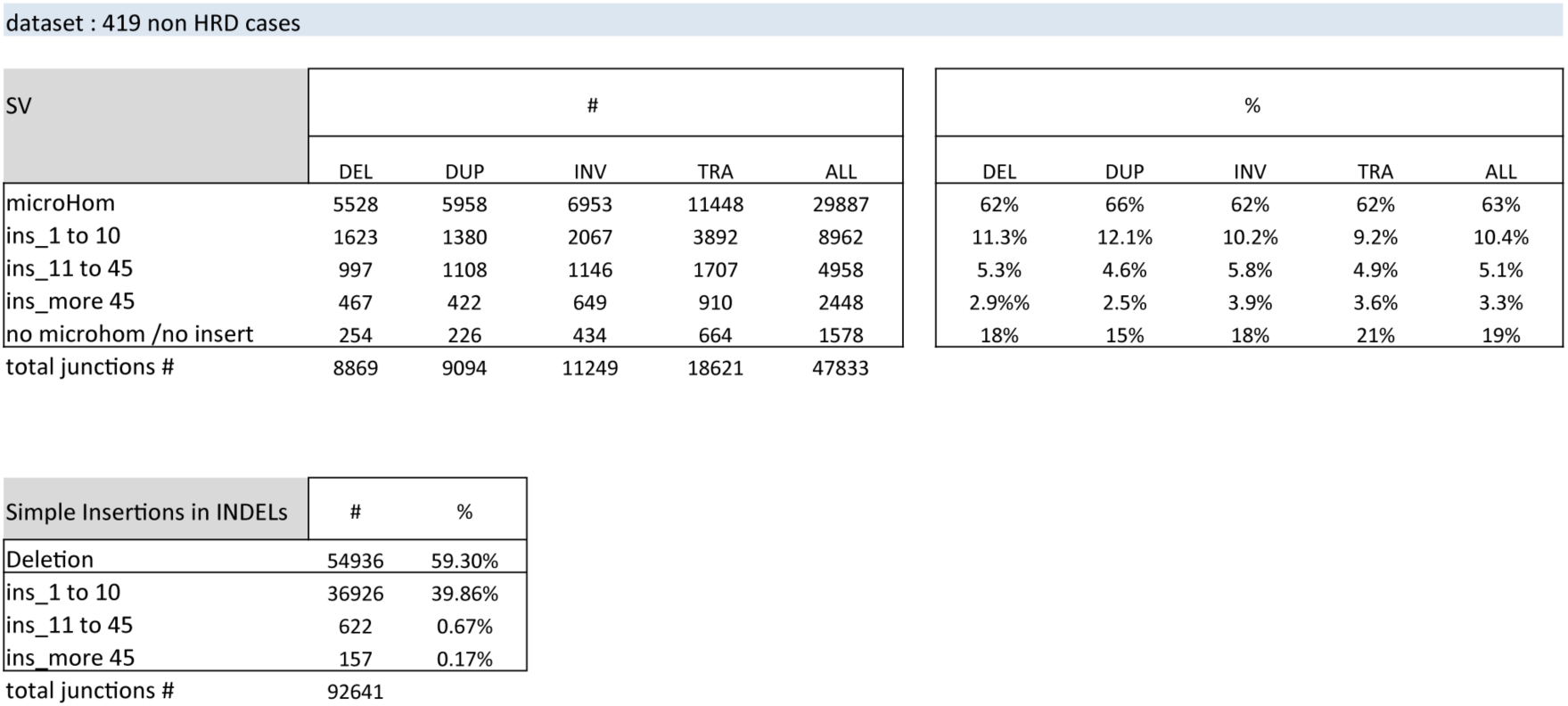
Analysis of SVs and INDELs junctions in non-HRD breast tumors.

